# UHMK1 is a novel splicing regulatory kinase

**DOI:** 10.1101/2022.05.21.492919

**Authors:** Vanessa C. Arfelli, Yun-Chien Chang, Johannes W. Bagnoli, Paul Kerbs, Felipe E. Ciamponi, Laissa M. S. Paz, Katlin B. Massirer, Wolfgang Enard, Bernhard Kuster, Philipp A. Greif, Leticia Fröhlich Archangelo

## Abstract

The U2AF Homology Motif Kinase 1 (UHMK1) is the only kinase that contains the U2AF homology motif (UHM), a common protein interaction domain among splicing factors. Through this motif, UHMK1 interacts with the splicing factors SF1 and SF3B1, known to participate in the 3’ splice site recognition during the early steps of spliceosome assembly. Although UHMK1 phosphorylates these splicing factors *in vitro*, the involvement of UHMK1 in RNA processing has not previously been demonstrated. Here, we identify novel putative substrates of this kinase and evaluate UHMK1 contribution to overall gene expression and splicing, by integrating global phosphoproteomics, RNA-seq, and bioinformatics approaches. Upon UHMK1 modulation, 163 unique phosphosites were differentially phosphorylated in 117 proteins, of which 106 are novel potential substrates of this kinase. Gene Ontology (GO) analysis showed enrichment of terms previously associated with UHMK1 function, such as mRNA splicing, cell cycle, cell division and microtubule organization. The majority of the annotated RNA-related proteins are components of the spliceosome, but are also involved in several steps of gene expression. Comprehensive analysis of splicing showed that UHMK1 affected over 200 alternative splicing events. Moreover, splicing reporter assay further supported UHMK1 function on splicing. Overall, RNA-seq data demonstrated that UHMK1 knockdown had a minor impact on gene expression and pointed to UHMK1 function in epithelial-mesenchymal transition. Finally, the functional assays demonstrated that UHMK1 modulation affects proliferation, colony formation, and migration of the cells. Taken together, our data implicate UHMK1 as a splicing regulatory kinase, connecting protein regulation through phosphorylation and gene expression in key cellular processes.

## Introduction

Phosphorylation is a reversible and dynamic post-translational modification mediated by kinases and counteracted by phosphatases. It is essential for the regulation of signaling networks that govern most of the cellular and physiologic processes, such as protein synthesis, cell growth and division, metabolism, inflammation, development, and aging [1]. The U2AF Homology Motif Kinase 1 (UHMK1) is the only kinase known to contain the U2AF homology motif (UHM) [2]. UHM motifs share a high level of sequence identity with the canonical RNA recognition motif (RRM), which mediates protein-RNA interactions and is often found in RNA-binding proteins. However, specific sequences within the UHM motif enable protein-protein instead of protein-RNA interactions. It has been proposed that UHMs evolved from RRMs and connect mRNA processing to other nuclear events [3]. UHMK1 is a serine/threonine kinase that preferentially phosphorylates proline-directed serine residues [4]. UHMK1 phosphorylates the splicing factors SF1 and SF3B1 *in vitro* [4,5] and interacts with these proteins through the UHM ligand motifs (ULMs) found in both factors [5]. SF1 is responsible for recognizing the branchpoint at the 3’ splice site of the introns during the early stages of spliceosome assembly [6] and participates in alternative splicing [7,8]. SF3B1 is a core component of U2 and U12 snRNP complexes required for canonical splicing [9]. The SPSP motif (S80 and S82) is the main phosphorylation site of SF1 and is targeted by UHMK1 *in vitro*. It has been suggested that phosphorylation of these residues possibly prevents premature association between the SF1-U2AF65 complex and the RNA [10]. Despite SF1 and SF3B phosphorylation being mediated by UHMK1, the direct involvement of this kinase in the splicing process has not been shown so far. UHMK1 also phosphorylates proteins involved in other cellular processes, such as cell cycle [11], cell migration [12], membrane trafficking [13], local translation in neurons [14], and cell differentiation [15,16]. In response to mitogens, UHMK1 is upregulated and phosphorylates the cyclin-dependent kinase inhibitor (CDKI) p27 on S10. This phosphorylation leads to p27 nuclear export and proteasomal degradation, counteracting the inhibitory effect of p27 on the cell cycle [11]. Similarly, UHMK1-mediated phosphorylation of Stathmin at S38 targets this protein to degradation, resulting in altered microtubule dynamics and impaired cell migration [12]. In neurons and endocrine cells, UHMK1 regulates vesicle secretion through phosphorylation of the Peptidylglycine α-amidating monooxygenase (PAM) cytosolic domain at S949, which is essential for the correct routing of PAM membranes [17–19]. Moreover, it was shown that UHMK1 enhances the local translation of β-actin and AMPA receptors, affecting spine morphology and post-synaptic activity of neurons [14,20]. Additionally, UHMK1 controls the differentiation of osteoclasts and osteoblasts [16], and is upregulated during the differentiation of hematopoietic cells [15]. More recently, UHMK1 was associated with cancer-related processes. In hepatocellular cancer, UHMK1 participates in the YAP-UHMK1-MYBL2 [21] and COX5B-UHMK1-ERK [22] axes, contributing to the malignant phenotype by altering the cell cycle and bioenergetics of liver tumor cells, respectively. By reprogramming the nucleotide metabolism through the UHMK1-NCOA3-ATF4 axis, UHMK1 contributes to gastric cancer [23]. Also, the involvement of UHMK1 in pancreatic cancer was reported [24].

To further characterize the UHMK1 function, we integrated quantitative global phosphoproteomics, RNA-seq, and splicing analysis using the vast-tools pipeline to identify novel putative substrates of this kinase and evaluate UHMK1 participation in gene expression and splicing. We identified a variety of novel UHMK1 candidate substrates that support the role of UHMK1 in biological processes previously associated with this kinase, particularly mRNA splicing. A comprehensive analysis of splicing and functional reporter assay showed for the first time that UHMK1 impacts mRNA splicing *in vivo*. Collectively, our data implicate UHMK1 as a splicing regulatory kinase, connecting protein regulation through phosphorylation and gene expression in key cellular processes.

## Results

### Global phosphoproteomic analysis revealed a range of putative substrates of the UHMK1 kinase

We performed untargeted quantitative phosphoproteomics of NIH3T3 cells depleted of UHMK1 (shUHMK1#2 and shUHMK1#3) and overexpressing the UHMK1 wild type (UHMK1^WT^) and the kinase-dead mutant (UHMK1^K54R^) (Figure 1a and b). We identified a total of 9861 phosphopeptides present in all three replicates of each sample, which were filtered by a p-value <0.05 and log2 fold change of ⍰1⍰ to identify differentially phosphorylated phosphopeptides (Figure 1c – f). In shUHMK1#2 and shUHMK1#3 knockdown cells, 12 and 85 phosphopeptides were differentially phosphorylated, respectively, compared to shCTRL control cells. In UHMK1^WT^ and UHMK1^K54R^ overexpressing cells, 53 and 37 phosphopeptides were differentially phosphorylated, respectively, compared to the empty vector (EV) control cells. The 187 differentially phosphorylated phosphopeptides encompassed 163 unique phosphosites within 117 unique differentially phosphorylated proteins (DPPs) (Figure 1g) regarded as UHMK1 putative substrates. Additionally, 25 out of the 187 differentially phosphorylated phosphopeptides contained multiple phosphosites whereas the remaining 162 phosphopeptides contained single phosphosites (Sup. Table 1). Among all phosphosites identified, 91% were serine, 8% threonine and 1% tyrosine residues (Figure 1h). Of note, 11 phosphosites (Sup. Table 1) were not registered in the PHOSIDA and PhosphositePlus databases, suggesting that they are specifically regulated upon UHMK1 modulation, as these phosphosites were not reported in other contexts.

**Figure 1.**
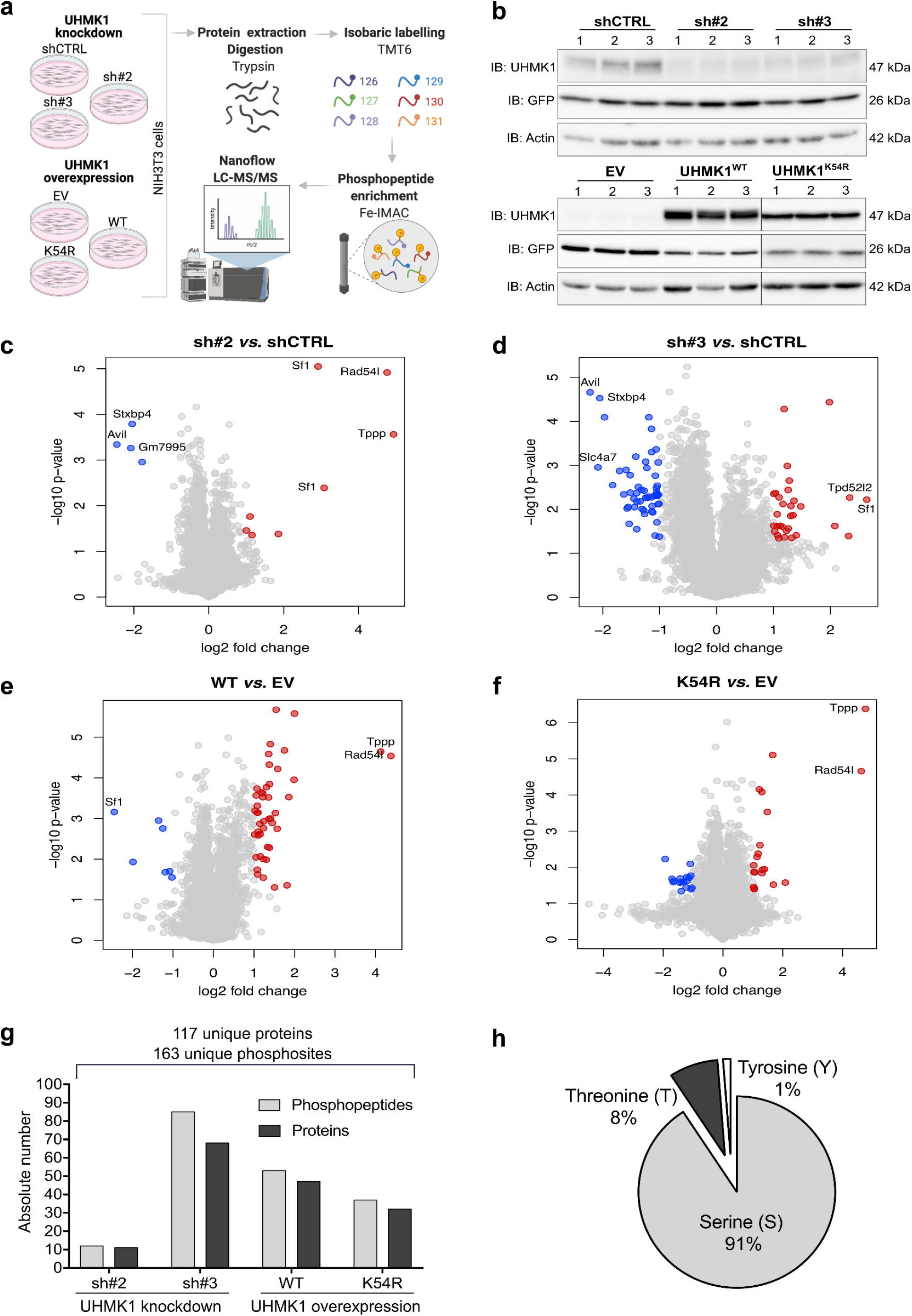
Global impact of UHMK1 on phosphoproteome. (a) Experimental workflow used in the phosphoproteome experiment. UHMK1 modulation in NIH3T3 cells was achieved expressing the shUHMK1#2 (sh#2) and shUHMK1#3 (sh#3) sequences, for knockdown, and UHMK1^WT^ (WT) or the kinase-dead mutant UHMK1^K54R^ (K54R) for overexpression. A scrambled shRNA sequence (shCTRL) and the empty vector (EV) MIY were used as control. Scheme created with ©BioRender.com. (b) Western blot confirming the efficient modulation of UHMK1 expression in NIH3T3 cells used in the phosphoproteome experiment. The numbers 1-3 represent the biological replicates from each condition. Membranes were blotted with anti-UHMK1, anti-GFP and anti-Actin (loading control). Total protein: 100 µg (knockdown) and 50 µg (overexpression). (c – f) The volcano plots show the global impact of UHMK1 knockdown (c,d) and overexpression (e,f) on phosphoproteome. The red and blue circles represent the significantly upregulated and downregulated phosphopeptides in each comparison, respectively. Phosphopeptides with p-value <0.01 and log2 fold change >|2|have their gene names assigned. (g) Overview of the absolute number of phosphopeptides and corresponding number of proteins differentially phosphorylated in the UHMK1 phosphoproteome. (h) Percentage of phosphorylated Serine, Threonine and Tyrosine residues in the UHMK1 phosphoproteome.

### Levels of UHMK1 kinase impact the phosphorylation of the identified substrates

Since virtually all DPPs in shUHMK1#2 were found in shUHMK1#3 cells (Sup. Figure 1) both lists were merged to comprise a total of 69 DPPs in UHMK1 knockdown cells (UHMK1-KD). The DPPs identified in UHMK1 knockdown and overexpressing cells (UHMK1^WT^ and UHMK1^K54R^) were compared in a Venn diagram (Figure 2a).

**Figure 2.**
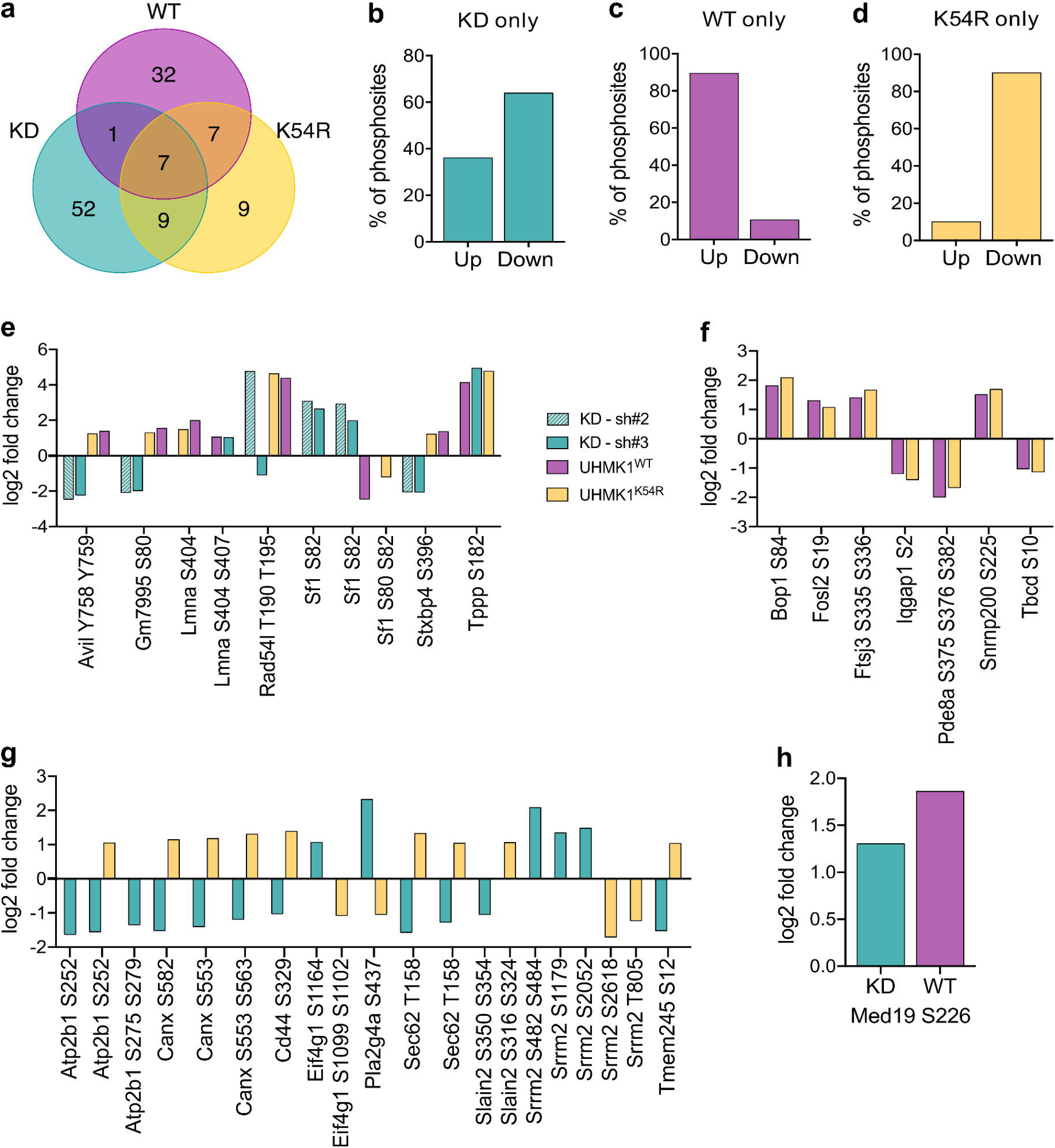
Distribution and phosphorylation pattern of the differentially phosphorylated proteins (DPPs). (a) Venn Diagram comparing the number of unique DPPs among the experimental conditions. KD = UHMK1-KD (knockdown); WT = UHMK1^WT^ overexpression; K54R= UHMK1^K54R^ overexpression. (b – d) Percentage of phosphosites up- and downregulated in the DPPs found exclusively in UHMK1-KD (b), UHMK1^WT^ (c), and UHMK1^K54R^ (d). (e – h) Phosphorylation pattern of the phosphopeptides (and respective phosphosites) from the DPPs (indicated by their gene names) shared by UHMK1-KD (sh#2 and sh#3), UHMK1^WT^ and UHMK1^K54R^ cells (e), UHMK1^WT^ and UHMK1^K54R^ cells (f), UHMK1-KD and UHMK1^K54R^ (g) and the only DPP (Med19) shared by UHMK1-KD and UHMK1^WT^ (h).

Several proteins were differentially phosphorylated exclusively in one condition, *i*.*e*., either in UHMK1-KD, UHMK1^WT^ or UHMK1^K54R^. Approximately 60% of phosphosites within the 52 proteins found exclusively in UHMK1-KD were downregulated, whereas the other 40% were upregulated (Figure 2b). Over 90% of the phosphosites within the 32 DPPs exclusively found in UHMK1^WT^ were upregulated (Figure 2c). Inversely, 80% of the phosphosites within the 9 proteins found solely in UHMK1^K54R^ were downregulated (Figure 2d).

Seven proteins, namely Advillin (Avil), Predicted gene 7995 (Gm7995), Prelamin-A/C (Lmna), DNA repair and recombination protein RAD54-like (Rad54l), Splicing factor 1 (SF1), Syntaxin-binding protein 4 (STXBP4) and Tubulin polymerization-promoting protein (TPPP), were differentially phosphorylated in all conditions (UHMK1-KD, UHMK1^WT^ and UHMK1^K54R^) (Figure 2e). Remarkably, the phosphorylation level of their phosphosites exhibited variable patterns among UHMK1-modulated conditions: the phosphosites Y758 and Y759 of Advillin, S396 of STXBP4 and S80 of Gm7995 were upregulated in UHMK1 overexpressing cells and downregulated in UHMK1-KD cells. Other phosphosites, such as S404 and S407 of Pre-lamin A/C and S182 of TPPP were upregulated in all conditions. Similarly, the phosphosites T190 and T195 of RAD54L were upregulated in the majority of the conditions, except in shUHMK1#3. Three phosphopeptides of SF1, a known UHMK1 substrate, were differentially phosphorylated upon UHMK1 modulation. Interestingly, two SF1 peptides comprised exclusively the phosphosite S82, which was downregulated in UHMK1^WT^ overexpression and upregulated in UHMK1 knockdown cells. Both S80 and S82 serines were differentially phosphorylated (downregulated) in a third phosphopeptide, but only in UHMK1^K54R^ overexpressing cells (Figure 2e).

The phosphosites of the DPPs in both UHMK1^WT^ and UHMK1^K54R^ exhibited the same phosphorylation pattern: either up or downregulated (Figure 2f). Conversely, the phosphosites of the 9 proteins regulated in both UHMK1-KD and UHMK1^K54R^ exhibited an opposite phosphorylation pattern in each condition (Figure 2g). The phosphosite S226 of Med19, the only protein regulated in UHMK1-KD and UHMK1^WT^, was upregulated in both conditions (Figure 2h).

Overall, these results show that UHMK1^WT^ overexpression leads to upregulation of the majority of the phosphosites, whereas the expression of the kinase-dead mutant (UHMK1^K54R^) or the UHMK1 knockdown leads to downregulation of most phosphosites. Thus, the phosphorylation pattern of the DPPs is strongly affected by the level of the kinase. Moreover, the data point to a dynamic and complex regulation of the differentially phosphorylated substrates present in more than one condition, some of which might be indirectly regulated by UHMK1.

### UHMK1 preferentially phosphorylates the ERXXSPEE consensus sequence

UHMK1 preferentially phosphorylates proline-directed serine residues *in vitro* [2]. We therefore sought to evaluate whether the same phosphorylation preference was present in our data. For these analyses, we considered the most likely direct targets of UHMK1, *i*.*e*., proteins whose phosphosites were upregulated in UHMK1^WT^ and downregulated in UHMK1-KD. Additionally, we included all the phosphosites regulated in UHMK1^K54R^, since many of these phosphosites were downregulated in this condition, having a similar effect of UHMK1-KD (Fig. 2d), or presented a pattern similar to UHMK1^WT^ (possibly due to residual endogenous activity, Fig. 2f and g). Using the software pLogo for motif search [25], we confirmed phospho-serines (S) followed by proline (P) at position +1 with high frequency (41.67%) and statistical significance (Figure 3a). Besides, glutamic acid (E) was present at positions -4 (17.71%), +2 (18.75%), and +3 (20.83%), while arginine (R) was common at position -3 in 16.67% of the phosphosites analyzed. As in our data the ERXXSPEE consensus sequence was not present in single phosphopeptides, this result indicates a preference of UHMK1 for these residues in these specific positions *in vivo*; however, the presence of the whole sequence is not a requirement for the phosphorylation of the serine residues. Additionally, UHMK1 also significantly phosphorylated proline-directed threonines (Figure 3b). However, the limited number of phospho-threonines matching the criteria used for this analysis (n=5) did not allow the identification of a broader consensus sequence. In summary, our data confirm UHMK1 as a preferential proline-directed kinase, with a preference for the amino acids ERXXSPEE surrounding the targeted serine residues.

**Figure 3.**
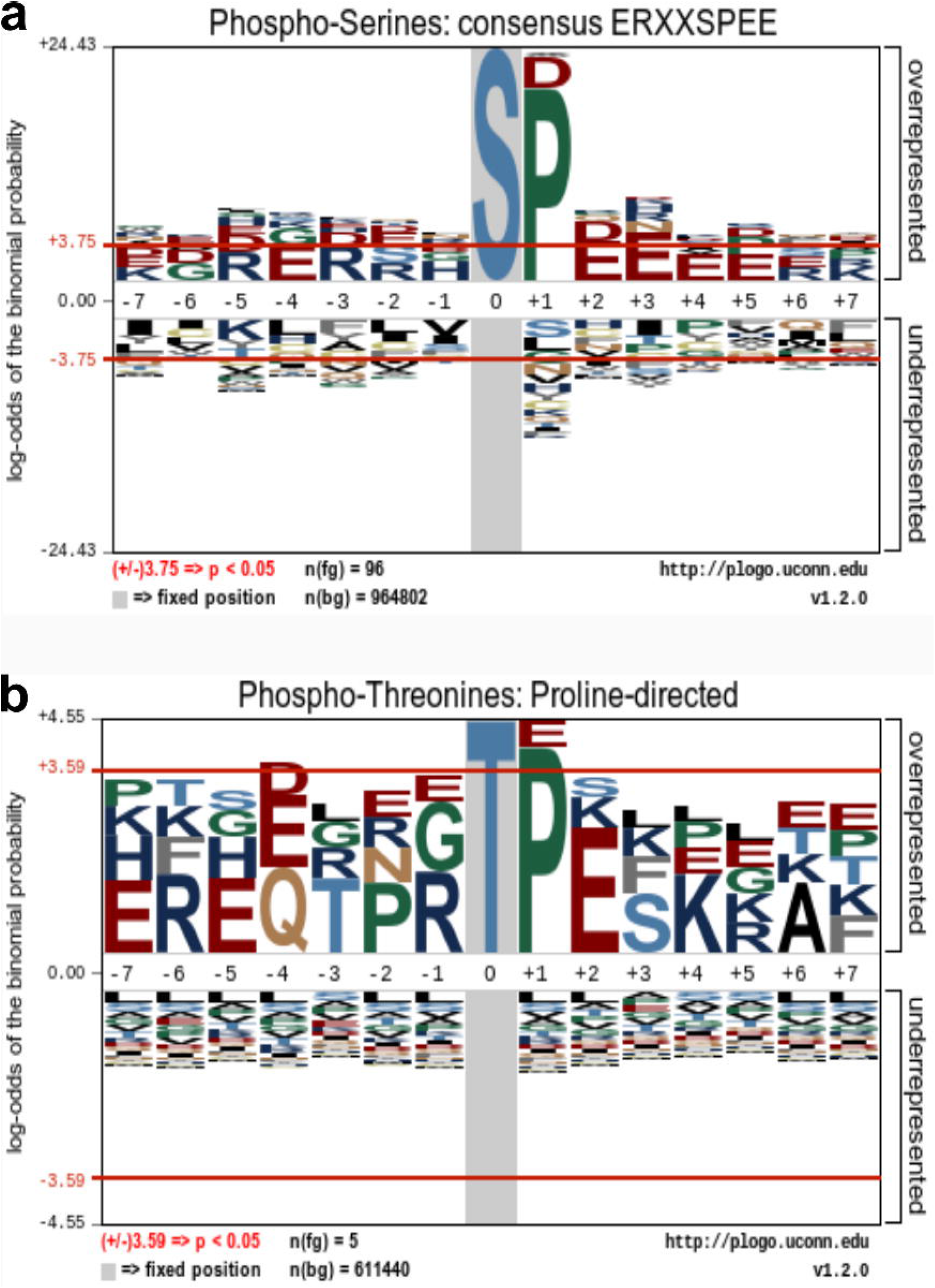
UHMK1 phosphorylation consensus sequence retrieved from pLogo analysis. Amino acid heights are scaled according to their statistical significance, as well as the stacking order (the most significant residues are positioned closest to the x axis). The red horizontal lines represent a threshold for Bonferroni-corrected statistical significance values. Positive values higher than the threshold correspond to statistically significant (p < 0.05) overrepresented amino acids. Negative values lower than the threshold correspond to statistically significant underrepresented amino acids. (a) UHMK1 preferentially phosphorylates proline-directed serine within the consensus sequence ERXXSPEE. Input sequences = 96. Log-odds of the binomial probability: E (-4) = 3.89; R (- 3) = 3.81; P (+1) = 20.32; E (+2) = 4.36; E (+3) = 5.07. (b) Proline-directed threonine is also preferentially phosphorylated by UHMK1. Input sequences = 5. P (+1) = 4.03.

### UHMK1 putative substrates are mainly implicated in RNA processing

To better understand the UHMK1 function, we performed a Gene Ontology (GO) enrichment analysis with the 117 DPPs regulated upon UHMK1 modulation. Among the Biological Process (BP) terms identified are “mRNA splicing via spliceosome”, “cell division”, “microtubule cytoskeleton organization”, “positive regulation of protein localization to cell periphery”, “translational initiation” and “maturation of LSU-rRNA from tricistronic rRNA transcript” (Figure 4a). Moreover, the terms “heterocyclic/aromatic compound metabolic process” and “nitrogen metabolic process” (and their related terms) recurrently appeared (Sup. Table 2). In addition, Reactome pathway analysis identified the pathways “eukaryotic translation elongation” (FDR= 0.00249) and “mRNA splicing – major pathway” (FDR= 0.00112) as statistically significant (not shown). The GO analysis confirmed the UHMK1 function in some cellular processes previously reported (cell cycle, microtubule cytoskeleton organization, translation, protein transport to the cell periphery) and previously suggested (mRNA splicing, cell division, nitrogen/aromatic compound metabolic process), as well as in a novel function, namely in rRNA processing.

**Figure 4.**
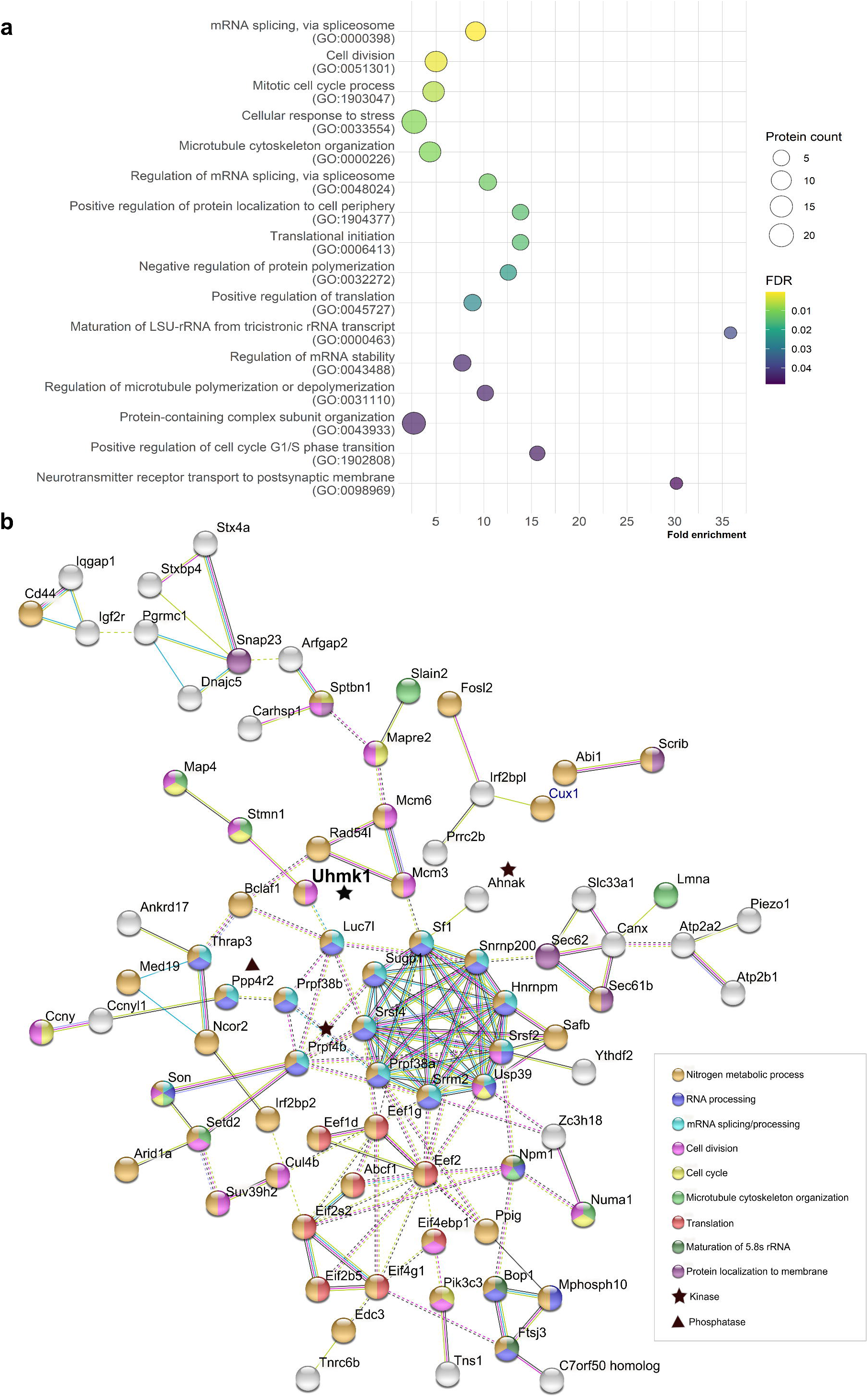
UHMK1 putative substrates are implicated in a variety of cellular processes and form a complex network. (a) Biological Process (BP) terms retrieved from Gene Ontology (GO) analysis. Only the most specific terms within the hierarchy presented in PANTHER are shown (complete results in Sup. Table 2, specific terms highlighted in grey). (b) Interaction network of UHMK1 and the 117 putative substrates (DPPs). Only the interacting proteins (represented by their gene names) are shown in the figure: the pink edges represent the experimentally determined interactions; blue edges represent interactions registered in curated databases; green edges represent interactions extracted from text mining; black edges represent co-expression. Each node represents one protein. Node colors indicate the most representative BP terms from GO analysis. Inter-cluster interactions are represented by dashed edges. Protein-protein interaction (PPI) enrichment p-value: 4.55e-15. The kinases and the phosphatase in this network are marked with a black star and a triangle, respectively. UHMK1 is highlighted in bold.

The interaction network of UHMK1 and the 117 putative substrates (DPPs) was accessed using the STRING database (Figure 4b). Proteins annotated in some of the BP terms retrieved from the GO analysis were identified as interaction clusters, from which the most prominent comprised the splicing regulatory proteins Sf1, Snrnp200, Hnrnpm, Srsf2, Usp39, Srrm2, Prpf38a, Srsf4, and Sugp1, together with Prpf4b, Prpf38b, Luc7l, Thrap3, Ppp4r2, and Son, which were presented as inter-cluster interactions. Of note, the kinases Ahnak and Prpf4b, and the phosphatase Ppp4r2 interact with proteins within this cluster, indicating an additional layer of regulation whereby UHMK1 could affect the phosphorylation of these targets. Moreover, there were marked intra-and inter-cluster interactions evident among translation factors (Eef2, Eef1d, Eef1g, Abcf1, Eif2s2, Eif2b5, Eif4g1, and Eif4ebp1). Yet, the proteins involved in rRNA processing (Bop, Mphosph10, Ftsj3, and Nmp1) and proteins that act in synaptic vesicles (Pgrmc1, Dnajc5, Snap23, Stx4a, and Stxbp4) formed two other additional clusters of interacting proteins. Interestingly, proteins related to cell cycle and cell division do not form an obvious interacting cluster but are also frequently associated with microtubule cytoskeleton organization (Numa1, Son, Stmn1, and Map4) (Figure 4b). In summary, our data demonstrate that UHMK1 regulates a large number of intrinsically connected RNA-related proteins.

### A subset of the RNA-related UHMK1 substrates contain putative ULM motifs

Since UHM domains are known to interact with UHM-ligand motif (ULM) of splicing factors [3], we searched for putative ULM motifs within the amino acid sequences of the 28 RNA-related DPPs annotated in the RNA-related BP terms of the GO analysis. Search of the [RK]-X(0,3)-W-[DN]-[EQ]ULM pattern [26] in ScanProsite returned two proteins owning *bona fide* ULM motif: SF1, whose ULM motif has been extensively characterized [5], and the RNA helicase TNRCB6. Since the ULM motif is highly degenerate and variations to the previously reported pattern may exist [26], we further investigated the putative ULM motifs in the 28 RNA-related proteins based on protein alignment. We first considered candidates for the alignment proteins bearing the conserved tryptophan (W) and at least two amino acids in assigned positions from the established ULM pattern. Those candidates were aligned with well-characterized ULM motifs from 4 other factors (U2AF^65^, SF3B1, ATX1, MAN1 [26]). Using this approach, we identified nine proteins with putative ULM motifs. Therefore, in addition to SF1, 10 out of the 28 RNA-related DPPs (TNR6CB, THRAP3, SNRNP200, EEF1D, SETD2, SAFB, SUGP1, USP39, FTSJ3, and PRPF38A) contain putative ULM domains (Figure 5) and are likely potential candidates for direct interaction with UHMK1.

**Figure 5.**
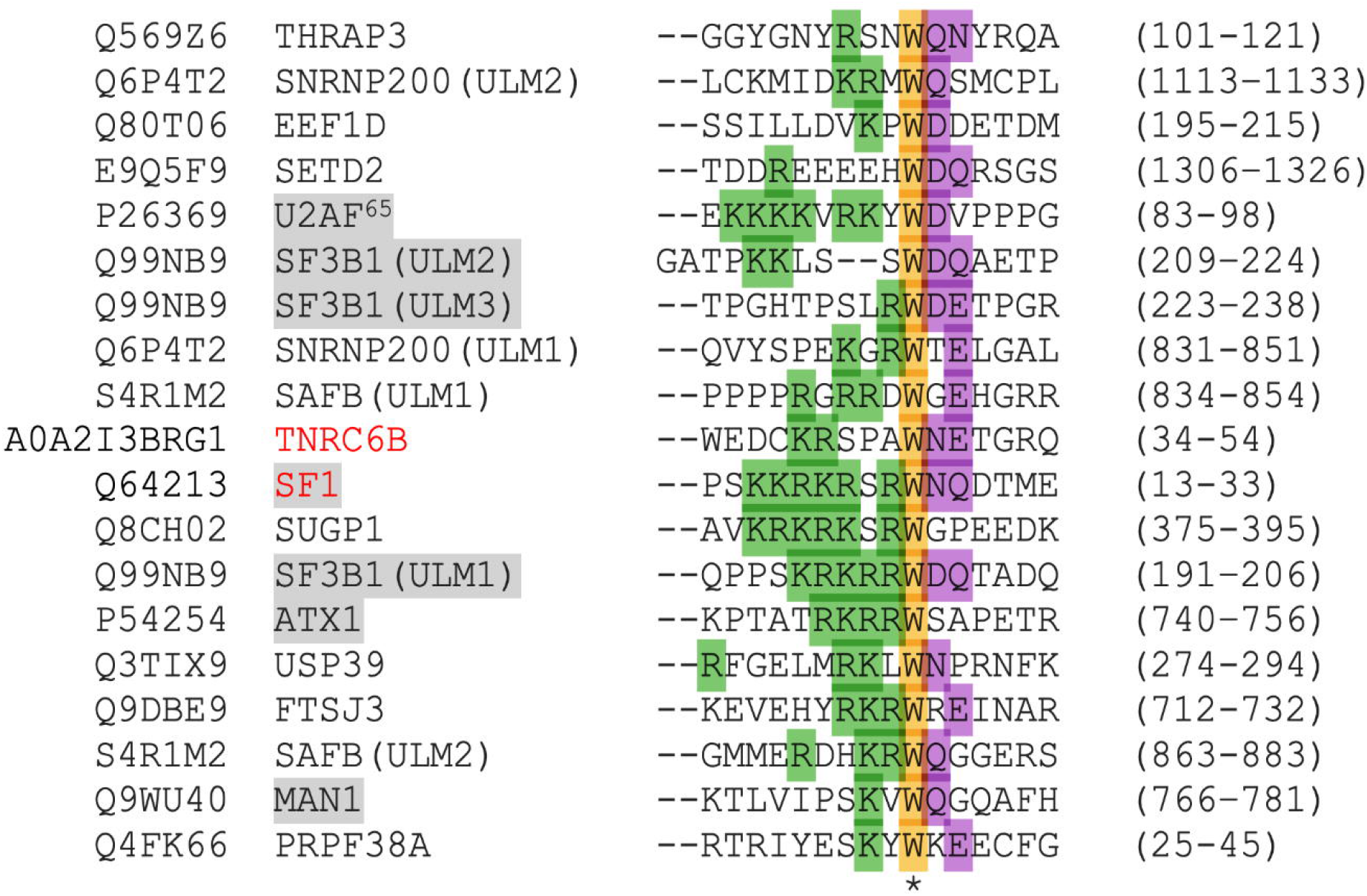
Besides SF1, 10 out of the 28 RNA-related DPPs contain ULM motifs. Clustal Omega alignment of proteins containing well characterized ULM motifs (highlighted in grey) and the 10 novel RNA-related putative UHMK1 substrates. Uniprot identification code for the mouse proteins are shown (left). The conserved tryptophan (W) within the ULM domain is highlighted in yellow. The preferred amino acids in the positions +1 and +2 are highlighted in purple, and the preferred amino acids at -1 and previous positions are highlighted in green. TNRC6B and SF1 (red) were the only RNA-related proteins from our study considered as *bona fide* ULM containing proteins in ScanProsite analysis (not shown).

### UHMK1 modulation impacts alternative splicing

Because several UHMK1 putative substrates are mainly related to mRNA splicing (Figure 4), we submitted UHMK1^WT^ overexpressing and knockdown NIH3T3 cells (Sup. Figure 2) to RNA-seq and investigated alternative splicing events (ASEs). A total of 179 statistically significant differentially spliced events were identified upon UHMK1 knockdown (Figure 6a) and 97 upon UHMK1^WT^ overexpression (Figure 6b). Overall, the most common type of event observed was the inclusion/exclusion of cassette exons, with 115 events (∼64% of total) occurring in UHMK1 knockdown and 51 events (∼52% of total) occurring after UHMK1^WT^ overexpression. Alternative 3’ splice sites (acceptor sites) and 5’ splice sites (donor sites) had a smaller sampling, with a total of 64 events in UHMK1 knockdown and 46 events in UHMK1^WT^ overexpression. No intronic events were observed in the conditions analyzed. Among all ASEs identified, 12 events were shared between UHMK1 overexpression and knockdown (Sup. Figure 3a), four of them (Dcun1d5:MmuALTD0004036-4/6, Cpsf6:MmuALTA0004602-4/4, Cpsf6:MmuALTA0004602-3/4, Cdca2:MmuALTA0003700-1/2) were altered in the same direction (included/excluded) in both experimental conditions, while the remaining eight events (Upk3bl:MmuALTA0000280-1/2, Upk3bl:MmuALTA0000280-2/2, Ktn1:MmuEX0025964, Mrps33:MmuALTD0008795-1/2, Mrps33:MmuALTD0008795-2/2, Insig2:MmuEX0024241, Safb2:MmuEX0040926, Kif20b:MmuEX0025516) were altered in opposite directions.

**Figure 6.**
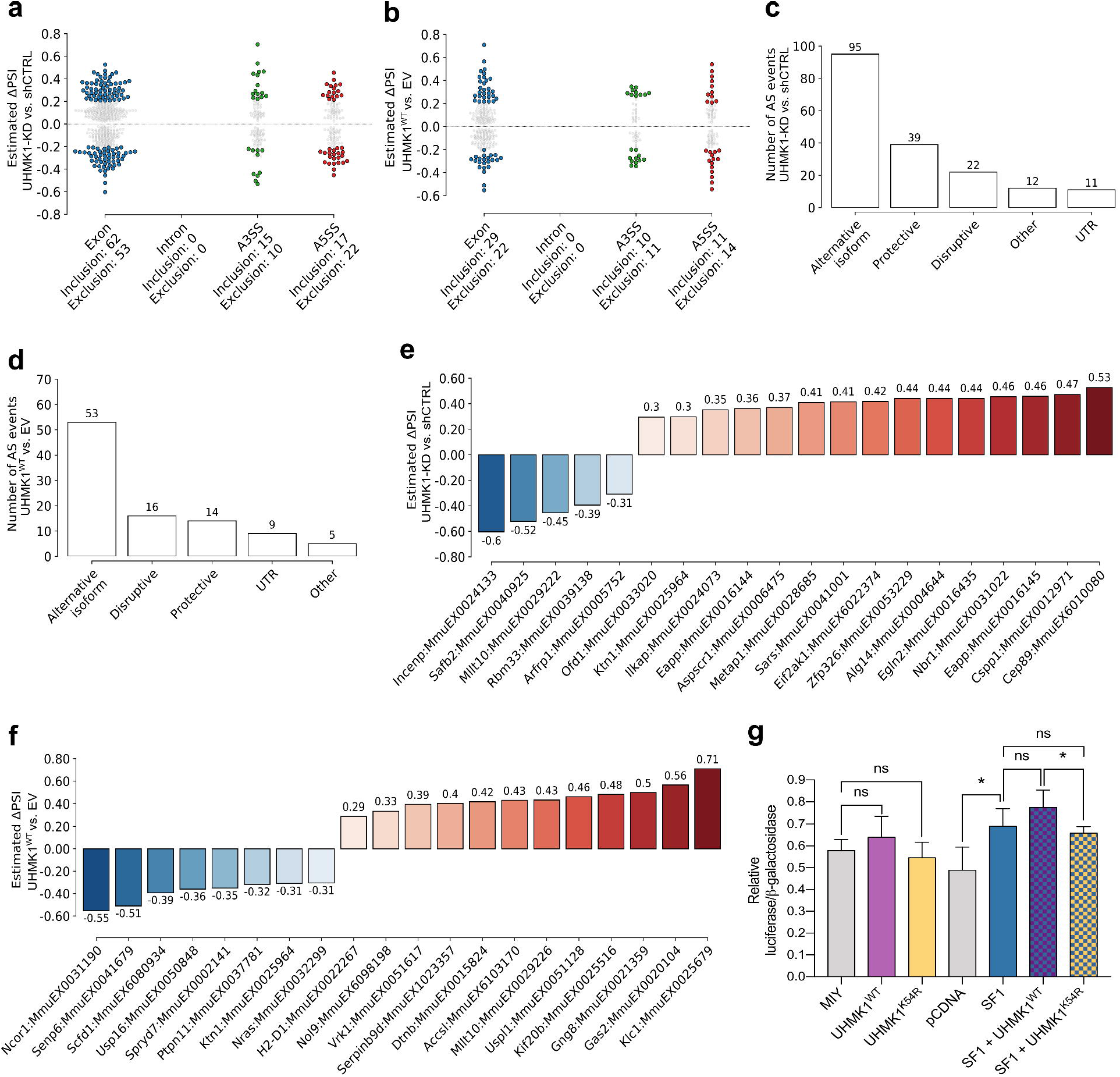
UHMK1 impacts mRNA splicing. (a – f) Splicing analysis from UHMK1 RNA-seq data, comparing UHMK1 knockdown (UHMK1-KD) with shCTRL cells, and UHMK1^WT^ overexpressing cells with the empty vector (EV) control cells. (a, b) Significant alternative splicing events (ASEs) in UHMK1-KD and UHMK1^WT^ overexpressing cells, respectively; (c, d) Predicted impact on coding sequence by ASE; (e, f) Percent spliced in (PSI) levels for the top 20 most significant Exon inclusion/exclusion ASEs. (g) Splicing reporter assay. The ratio between β-galactosidase and luciferase is an indirect measure of splicing occurring between both genes. Average values of four independent experiments. Constant amount of the pTN24 reporter plasmid is present in all conditions. One way ANOVA followed by Bonferroni correction was used to compare UHMK1^WT^ and UHMK1^K54R^ to the empty vector control MIY (not significant). Increase in splicing in SF1 over pcDNA (p=0.0244) and difference between SF1+UHMK1^WT^ and SF1+UHMK1^K54R^ (p=0.0294) was confirmed by Student T test. Controls of gene expression levels of UHMK1 and SF1 are provided in Sup. Figure 4.

Analysis of the predicted impact of the ASEs on coding sequences revealed that 53% (95 events) of ASEs in UHMK1 knockdown cells mainly predicted alternative isoforms with neutral impact, whereas 12% (22 events) exhibited a disruptive, and 21 % (39 events) a protective impact on the ORF of the proteins they encode (Figure 6c). In UHMK1^WT^ overexpressing cells, 54% (54 events) of the splicing changes were predicted as alternative isoform (neutral), 16% (16 events) were predicted to disrupt the ORF and 14% (14 events) predicted a protective function (Figure 6d). The percent spliced in (PSI) of the 20 most significant cassette exons are depicted in Figure 6e and f. Among those, disruptive ASEs were found in the transcripts of *Safb2, Mllt10*, and *Ktn1* (UHMK1 knockdown) and *Usp16, Ptpn11, Scfd1*, and *Dtnb* (UHMK1^WT^). Interestingly, GO analysis of the list of genes exhibiting ASEs in UHMK1 knockdown returned BP terms mainly related to cell-cycle, microtubule organization and metabolic process (Sup. Figure 3b), which were also found in the GO analysis of the UHMK1 phosphoproteome (Figure 4).

To further explore the contributions of UHMK1 to splicing regulation, we performed an *in vivo* splicing reporter assay. HEK 293T cells were transfected with the reporter plasmid and UHMK1^WT^ or UHMK1^K54R^ alone and in combination with the UHMK1 substrate, SF1. Expression of UHMK1^WT^ alone increased the activity of the reporter only by 10% over the control (empty vector MIY), whereas splicing activity was barely observed upon expression of the UHMK1 kinase dead-mutant (-6% compared to the control) (Figure 6g). To evaluate the impact of UHMK1 on SF1-mediated splicing, UHMK1 was co-expressed with SF1 and the reporter plasmid. Expression of SF1 alone enhanced the activity of the reporter by 41 % over the control (empty vector pcDNA). Co-expression of UHMK1^WT^ enhanced SF1 function on the splicing of the reporter gene by 18% compared to SF1 alone. The effect of UHMK1 on SF1 was dependent on the kinase activity since expression of the UHMK1^K54R^ did not enhance SF1-mediated splicing (-25% compared to SF1+UHMK1^WT^, p=0.0294). Although the differences were subtle, they were consistently observed in 4 independent experiments. Levels of *UHMK1* and *SF1* expression are shown in Sup. Figure 4. Taken together, our data show for the first time that UHMK1 modulation can impact splicing and implicate UHMK1 as a regulatory splicing kinase.

### UHMK1 has a modest impact on global gene expression

Since splicing and gene expression are tightly coordinated processes [27] we sought to evaluate the UHMK1 impact on gene expression by RNA-seq. Differential expression analysis revealed that modulation of UHMK1 had a modest impact on global gene expression, as observed in the Principal Component Analysis (PCA) plots (Figure 7a and b). UHMK1 knockdown (UHMK1-KD), mediated by the expression of shUHMK1#1, shUHMK1#2, and shUHMK1#3 sequences, altered the expression of 32 genes, of which, 17 were upregulated and 15 were downregulated (Fig. 7c and d, and Sup. Table 3). UHMK1^WT^ overexpression did not affect gene expression, as *Uhmk1* was the only differentially expressed gene in this setting (data not shown). Although UHMK1 knockdown had a small impact on expression, with log2 fold changes < ⍰2⍰ (Sup. Table 3), we could validate these findings by a qPCR array of selected targets in an independent transduction experiment of NIH3T3 cells (Sup. Figure 5).

**Figure 7.**
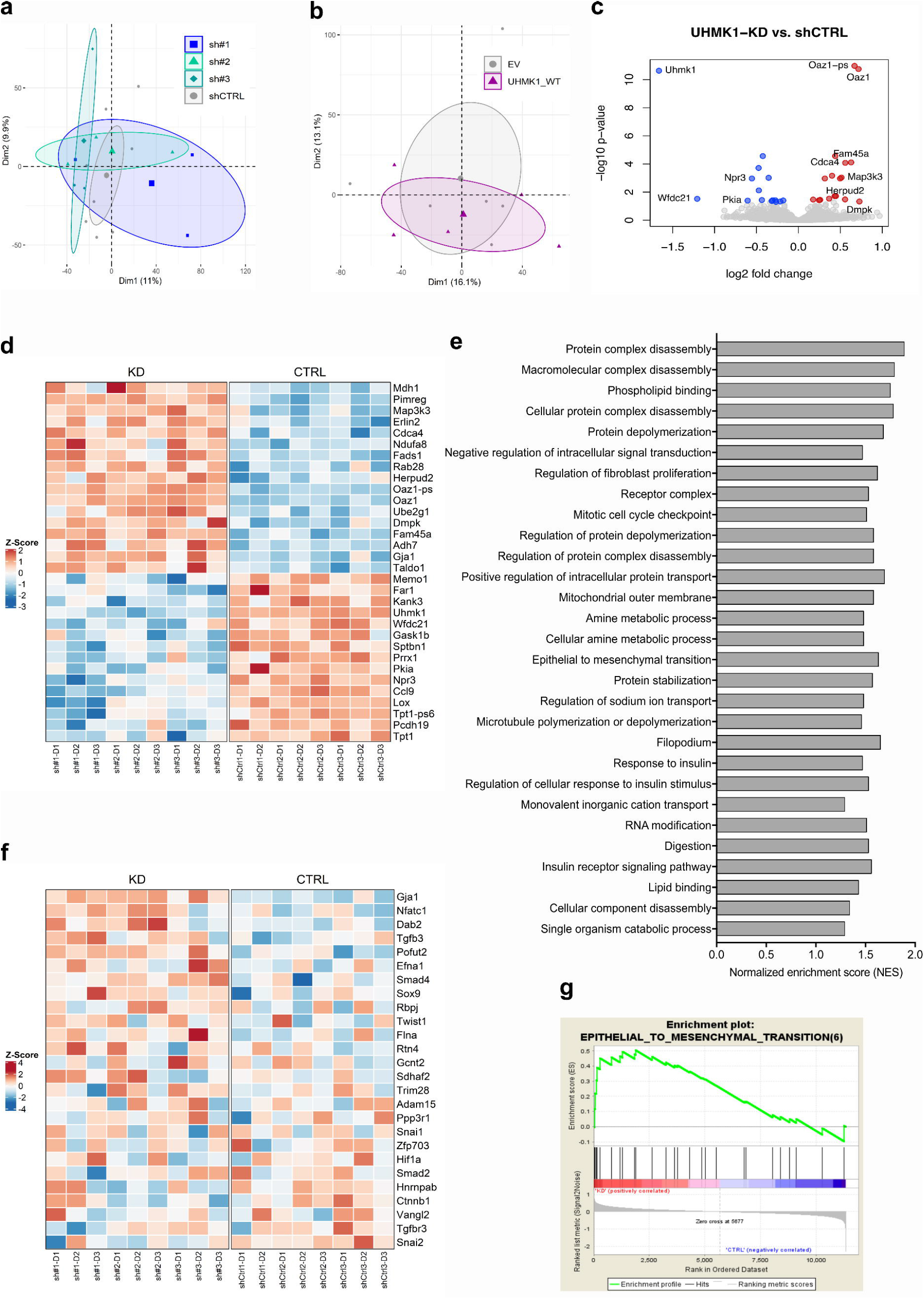
UHMK1 impacts gene expression. (a, b) Principal Component Analysis (PCA) plots showing the overall effect of UHMK1 knockdown mediated by shUHMK1#1 (sh#1), shUHMK1#2 (sh#2) and shUHMK1#3 (sh#3) compared to shCTRL, and UHMK1^WT^ overexpression compared to empty vector (EV) control, respectively. (c) Volcano plot showing the differentially expressed genes (DEGs) in UHMK1-KD cells (shUHMK1#1, shUHMK1#2 and shUHMK1#3 combined as one single artificial condition) compared to shCTRL cells. p-value < 0.05 was used as cutoff. Genes with log2 fold change > |0.5|are labeled. (d) Heatmap of expression of the 32 DEGs represented as z-score of normalized counts, in each replicate of UHMK1 knockdown (KD) and shCTRL (CTRL) cells. (e) Gene sets enriched in UHMK1-KD in GSEA analysis. (f) Heatmap showing the expression of the EMT enriched gene set in UHMK1 knockdown cells. Gene expression is represented as z-score of normalized counts. (g) Enrichment plot for Epithelial Mesenchymal Transition (EMT) gene signature.

### 2GSEA analysis implicates UHMK1 in epithelial-mesenchymal transition

Gene Set Enrichment Analysis (GSEA) revealed 29 enriched gene sets in UHMK1 knockdown cells, considering a nominal p-value < 0.05 (Figure 7e). Although none of the gene sets crossed the 25% false discovery rate (FDR) threshold (<0.25), 4 out of 29 were related to functions previously associated with UHMK1 in the literature, namely “regulation of fibroblast proliferation” and “mitotic cell cycle checkpoint”; “microtubule polymerization or depolymerization”, and “RNA modification”. Moreover, these 4 gene sets were also observed as related terms in GO analysis of the UHMK1 regulated proteins (Figure 4 and Sup. Table 2). Therefore, it is plausible that the 29 enriched gene sets are true UHMK1-related pathways. Among the enriched gene sets, epithelial-mesenchymal transition (EMT) showed the highest percentage of genes with dichotomized expression pattern between UHMK1 knockdown and control samples (Figure 7f and g) suggesting that UHMK1 may play a role in EMT.

### UHMK1 affects proliferation, clonogenicity, and migration of NIH3T3 cells

Because the GO and GSEA analysis returned terms related to cell cycle, cell division, and migration, we performed functional assays with UHMK1 overexpressing and knockdown NIH3T3 cells to evaluate the phenotypes related to those cellular processes (Figure 8). Proliferation was significantly reduced in UHMK1 knockdown cells (shUHMK1#1 and shUHMK1#2), whereas in UHMK1^WT^ overexpressing cells there were no effects on proliferation (Figure 8a). In colony-forming assay, no difference in the percentage of colonies formed was observed with shUHMK1#1, shUHMK1#2 or shUHMK1#3 cells compared to shCTRL cells, while UHMK1^WT^ overexpressing cells formed 20% less colonies than control cells (Figure 8b). Viability and apoptosis were also not affected upon UHMK1 modulation (Figure 8c and d).

**Figure 8.**
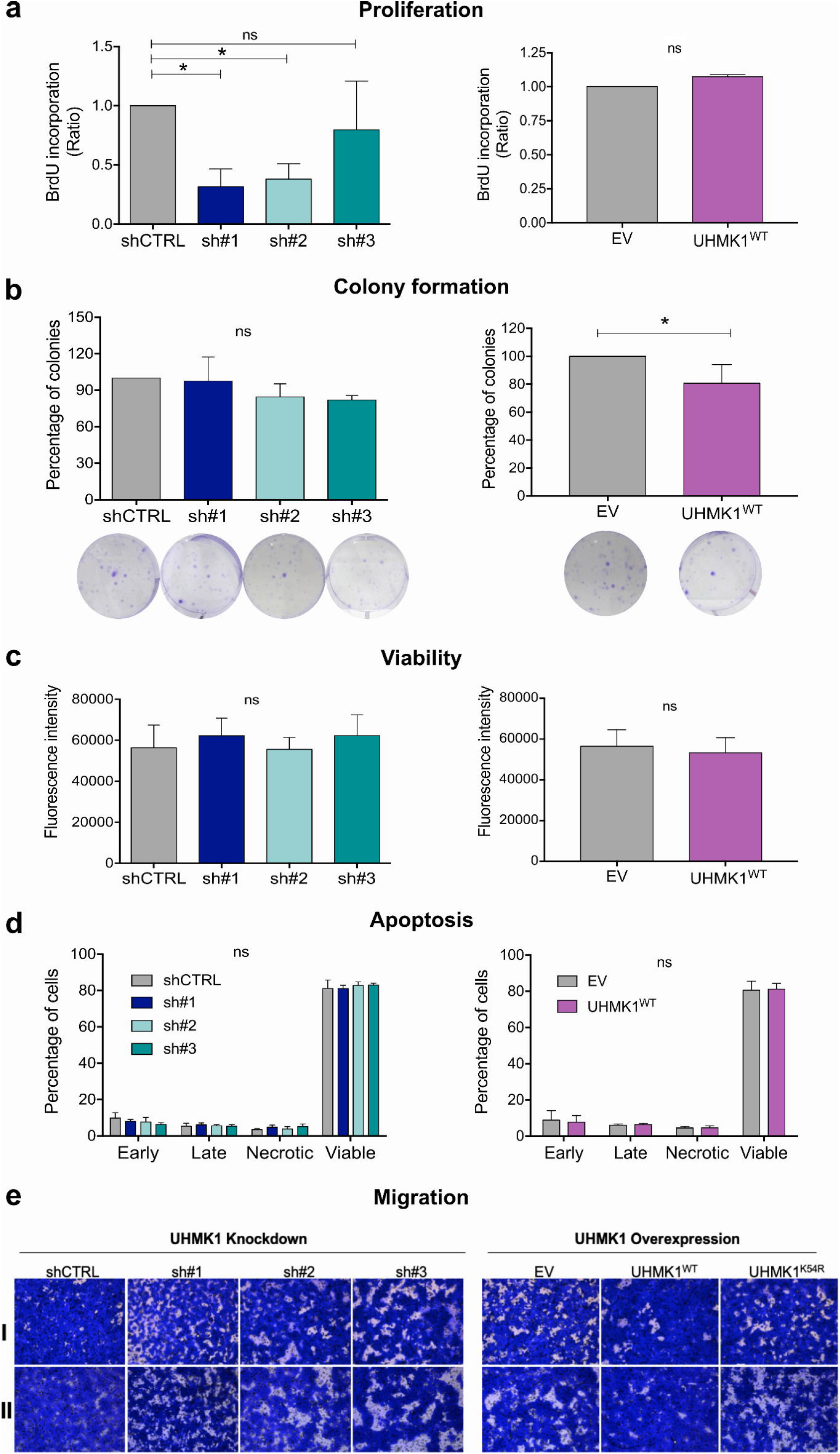
UHMK1 affects proliferation, colony formation, and migration of NIH3T3 cells. (a) Proliferation was evaluated by the percentage of cells that incorporated BrdU. Charts represent the ratio of BrdU incorporation in UHMK1 knockdown mediated by shUHMK1#1 (sh#1), shUHMK1#2 (sh#2) and shUHMK1#3 (sh#3) relative to the scrambled control (shCTRL) cells, and UHMK1^WT^ overexpressing cells relative to empty vector (EV) cells. Data from three independent experiments (* = p < 0.05, One-way ANOVA followed by Bonferroni’s multiple comparison test). (b) Clonogenic assay. The bar plots represent the percentage of colonies in UHMK1 knockdown and UHMK1^WT^ overexpressing cells relative to the shCTRL or empty vector (EV), respectively. Results from three independent experiments, carried out in triplicate (* p= 0.0286, Mann-Whitney test). The representative images of crystal violet-stained colonies are shown. One colony was defined by the minimum of 50 cells. (c) Viability assay. For UHMK1 knockdown cells, the chart represents the average of four experiments performed in sextuplicate. For UHMK1^WT^ overexpressing cells, the represents the average of 5 experiments, 2 performed in triplicate and 3 performed in sextuplicate. (d) Apoptosis evaluation by Annexin V assay. Mean from 3 independent experiments. Early apoptosis is defined by the cells that are positive only for Annexin V; Late apoptosis comprises the cells that are positive for both Annexin V and propidium iodide (PI); Necrotic cells are positive only for PI and Viable cells are negative for both markers. (e) Migration assay. Representative imagens of two independent experiments (I and II) with UHMK1 knockdown and UHMK1^WT^ overexpressing cells. The experiments were performed in duplicates. 3 images from each replicate were taken: in the center, bottom and top of the membrane, relative to position of the circumference in the plate. The images in this figure are representative of the center. The control representing spontaneous migration towards lower chamber containing 1% FBS, and the images acquired in the top and bottom of the wells are provided Sup. Figure 6, 7 and 8. Images were acquired with Microscope Leica DMi8, 10 x magnification.

Finally, transwell chemotaxis assay revealed that UHMK1 knockdown cells migrated less than shCTRL cells (Figure 8e left panel and Sup. Figure 6 and 7). Conversely, UHMK1^WT^ overexpression increased migration of cells towards the gradient, whereas overexpression of the UHMK1 kinase-dead mutant (UHMK1^K54R^) had no effect on cell migration and was comparable to control cells (Figure 8e right panel and Sup. Figure 6 and 8).

## DISCUSSION

The implication of UHMK1 function in cellular processes such as cell cycle, migration, membrane trafficking, local translation in neurons, and mRNA metabolism is mainly based on the knowledge of the UHMK1 interaction with particular binding partners involved in these processes [4,11,12,14,19]. To our knowledge, this study is the first large-scale investigation of the UHMK1 phosphoproteome, carried out using UHMK1-depleted and overexpressing cells.

Our phosphoproteome results showed that UHMK1 regulates phosphosites of 117 proteins, of which 106 are novel putative substrates for the kinase. The fact that 10% of the proteins identified are either known UHMK1 substrates, such as SF1 [4], Stathmin [12,28], and NPM1 [22]; or have been previously described as potential UHMK1 interacting proteins, such as STXBP4, TNRC6B, IRF2BP2, NCOR2, YTHDF2, SUGP1, hnRNPM, and PRRC2B [21], points to the accuracy of our findings. Even though some of the UHMK1 known substrates did not appear in this initial screen, for instance, SF3B1 [5], p27 [11], and PIMREG [29], they were identified in one or other experimental conditions below the cut off employed, further confirming the stringency of the study. Of note, five kinases (PIK3C3, WNK1, NUCKS1, PRPF4B, and AHNAK) and two phosphatases (PPP4R2 and PTPN21) were found among the UHMK1 putative substrates. These proteins likely contribute to additional layers of regulation of the DPPs, some of which might be only indirectly regulated by UHMK1, through these kinases and phosphatases.

Gene Ontology enrichment analysis of the 117 putative substrates confirmed UHMK1 function in cellular processes previously associated with this kinase (mRNA splicing [4], cell cycle [11]; cytoskeleton (microtubule) organization [12]; translation [14,20], membrane trafficking [18,19], and nucleotide metabolism [23]) and pointed to a novel function of UHMK1 in rRNA processing. Most importantly, our data broadened the knowledge of the protein network and the players regulated by UHMK1 in each of these processes. Moreover, the fact that many of the UHMK1 putative substrates interact with each other and between the clusters (Figure 4b) indicates that UHMK1 is important for regulating complex networks of proteins that coordinate diverse (and yet complementary) biological processes in the cell. Remarkably, 24% of the putative UHMK1 substrates are RNA-related proteins and the most prominent interaction network in STRING analysis is comprised of splicing regulatory factors, followed by translation factors, and rRNA processing proteins. A closer look at the function of the most prominent group revealed that most of the proteins are involved in the spliceosome, at the complexes E, A, and B, but also in the further steps of the splicing cycle (Figure 9). It is well known that the phosphorylation status of splicing factors impacts the assembly of the spliceosome, as it affects interaction among them and with the mRNA. Moreover, phosphorylation controls the intracellular and intranuclear localization of splicing factors, altering their concentration and consequently splice site selection [30]. Two major families of splicing regulatory kinases, namely the SRKPs and CLKs are responsible for phosphorylating splicing factors [31]. This is the first report describing UHMK1 as a broad splicing regulatory kinase. However, the consequences of the UHMK1-mediated regulation of the phosphosites of these proteins remain to be investigated.

**Figure 9.**
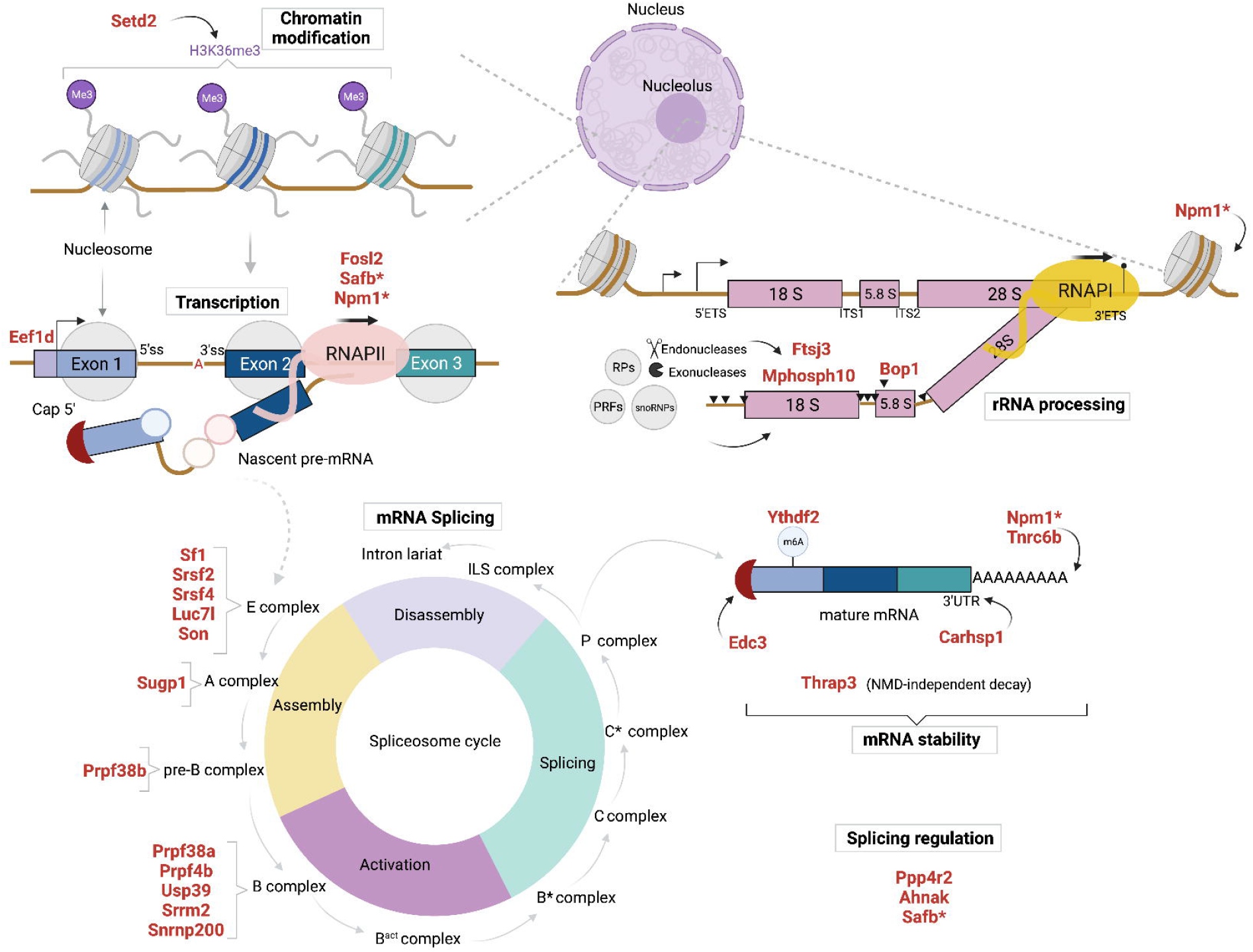
Schematic view of the role of the 28 RNA-related UHMK1 substrates (DPPs) in different layers of gene expression. We searched the role of the 28 RNA-related DPPs in the literature, UniProt [80] and Spliceosome [81] databases. The RNA-related DPPs are highlighted in red. Proteins that act in more than one function related to RNA metabolism are marked with an asterisk (*). The main functions are highlighted in a box: Chromatin modification, Transcription, mRNA splicing, Splicing regulation, mRNA stability and rRNA processing. RPs: ribosomal proteins; PRFs: pre-ribosomal factors. Approximately half of the novel RNA-related UHMK1 substrates (15 proteins) are components of the spliceosome. Six DPPs have a role in mRNA stability, through mechanisms involving the poly(A) tail (Tnrc6b [82] and Npm1 [83]), the 5’ Cap (Edc3 [84]), binding to N6-methyladenosine mRNAs (Ythdf2 [85]), nonsense-mediated decay (NMD)-independent decay (Thrap3 [86]), and binding to 3’ UTR (Carhsp1 [87]). Four of the RNA-related DPPs act in transcription, either by acting on RNA pol II regulation (Safb [88] and Npm1 [89]) or as transcription factors (Fosl2 [90] and Eef1d [91]). Setd2, a methyltransferase responsible for H3K36me3 mark, act in chromatin modification [92]. Moreover, three DPPS act specifically in the cleavage of the tricistronic rRNA transcript: Mphosph10 (component of the U3 snoRNP) and Ftsj3 (2’-O-methyltransferase) are involved in the processing of the 18S rRNA [93,94], while Bop1 (component of the PeBoW complex) is necessary for the processing of 5.8S and 28S rRNAs [95]. Npm1 also acts in the nucleolus as a histone chaperone, affecting chromatin status of rDNA [89]. Created with ©BioRender.com.

Splicing analysis of transcriptome data revealed that UHMK1 modulation affected over 200 alternative splicing events, most of which led to exon inclusion/exclusion and produced alternative isoforms. Since our analysis was performed on Prime-seq data [32] and the reads were sequenced mainly from the 3’ end of the transcripts, one limitation of our analysis is that we only evaluated alternative splicing events occurring within this region of the transcripts. Thus, we cannot consider it a global splicing analysis as the identified ASEs are likely to be underrepresented. Nonetheless, our data is a proof-of-principle that UHMK1 affects alternative splicing and it applies to many target genes. A reporter assay confirmed that UHMK1 influences splicing *in vivo* and showed that this effect is dependent on the UHMK1 kinase activity. These data, together with the knowledge that UHMK1 regulates the phosphorylation of a complex network of splicing factors, suggest that UHMK1 function on splicing is more indirect, through the action of the various splicing related factors as well as kinases (AHNAK, PRPF4B) and phosphatases (PPP4R2) regulated by UHMK1.

We showed that the expression of ectopic UHMK1^WT^ was able to increase the SF1-mediated splicing of a reporter gene. SF1 is a well-known substrate of UHMK1 *in vitro* [4,5,10,33]. We identified SF1 as the most significant differentially phosphorylated protein in our phosphoproteome analysis, confirming that UHMK1 controls the phosphorylation status of the SF1 S80 S82 residues (SPSP motif) *in vivo*. However, our observation that only S82 was significantly upregulated in UHMK1-KD, and downregulated in UHMK1^WT^ overexpression, while both S80 and S82 were significantly downregulated in UHMK1^K54R^ is quite puzzling and suggests that a complex and rather indirect regulation of SF1 by UHMK1 *in vivo* is likely. Supporting this idea, the expression of UHMK1 was unable to increase the incorporation of radioactive phosphate on full-length SF1 in mammalian cells [4]. Moreover, the SF1 SPSP motif is normally found in a highly phosphorylated state in proliferating cells [33], and a subset of SF1 remains phosphorylated in UHMK1 knockout mice [34]. Besides UHMK1, SRPK2 is also known to phosphorylate the SF1 SPSP motif, preferentially at the residue S82 [35]. Thus, the SF1 SPSP motif is likely to be regulated by different kinases and phosphatases [10]. Interestingly, the Serine/threonine-protein phosphatase 4 regulatory subunit 2 (Ppp4r2) was found upregulated in UHMK1^WT^ expressing cells and the Serine/threonine-protein kinase PRP4 homolog (Prpf4b) was found upregulated in UHMK1-KD cells, making it tempting to speculate that these enzymes could be responsible for the SF1 SPSP dephosphorylation and phosphorylation observed in these respective conditions. Yet, we cannot rule out that the reversed phosphorylation pattern observed on SF1 SPSP motif was due to the cell context and/or prolonged exposure to UHMK1 modulation in our experimental conditions. Altogether, we showed that UHMK1 is the kinase controlling the phosphorylation status of SF1 SPSP motif *in vivo* and that the regulatory mechanism is complex and might involve additional players other than a direct UHMK1 targeted substrate regulation as previously described [4].

It is well accepted that mRNA splicing and transcription are coupled events, and as such, one affects the other [36]. Similarly, capping, cleavage and polyadenylation occur co-transcriptionally. Moreover, the chromatin state and the type of promoter influence the recruitment of transcription and splicing factors, as well as the kinetics of RNA pol II, which in turn influences mRNA transcription rate and alternative splicing [37]. Indeed, the RNA-related proteins found in our phosphoproteome data are involved in several steps controlling gene expression, from chromatin modification to mRNA splicing and cleavage to generate mature rRNA (Figure 9).

Splicing-associated chromatin signatures (SACS), characterized by the combination of specific histone marks, have been recently described as a mechanism to rapidly adjust alternative splicing [38]. Here, we found that the phosphorylation on S743 of SETD2, a histone lysine methyltransferase responsible for H3 lysine 36 trimethylation (H3K36me3), is upregulated in UHMK1^WT^. Moreover, a predicted protective ASE transcript of *Setd2* was identified in UHMK1-KD (not shown). Besides SETD2, other chromatin modifier enzymes (SUV39H2, L3MBTL2, NCOR2, ARID1A, CUL4B) were found differentially phosphorylated in our study.

Finally, the UHMK1-mediated regulation of proteins involved in rRNA processing, namely pre-rRNA 2’-O-ribose RNA methyltransferase FTSJ3 (Ftsj3), Ribosome biogenesis protein BOP1 (Bop1), Nucleophosmin (Npm1), and U3 small nucleolar ribonucleoprotein protein MPP10 (Mphosph10) is a new finding. This points to a specific role of UHMK1 in the control of rRNA processing and ribosome biogenesis. These processes are closely associated with protein translation [39]. In fact, UHMK1 regulated the phosphorylation of 8 translation factors: EF-2 (Eef2), Translation initiation factor eIF-2B subunit epsilon (Eif2b5), eIF-2-beta (Eif2s2), EF-1-delta (Eef1d), eEF-1B gamma (Eef1g), eIF4E-binding protein 1 (Eif4ebp1), eIF-4-gamma 1 (Eif4g1), and ATP-binding cassette sub-family F member 1 (Abcf1). Taken together, our results show that UHMK1 controls phosphorylation of proteins involved in different steps of gene expression regulation. This is in agreement with our previous report where we showed that UHMK1 could influence the expression of a report gene [29] and with other studies showing the effect of UHMK1 on gene expression [13,21,34].

Intriguingly, in our RNA-seq analysis, UHMK1 knockdown had limited impact on gene expression, with the identification of 32 DEGs. Nevertheless, some of the gene signatures retrieved from the GSEA analysis relate to the biological processes identified in the GO analysis of phosphoproteome data (Protein complex assembly/disassembly, Microtubule and cytoskeleton organization, Amine/nitrogen metabolic processes, RNA modification, Cell-cycle and Cell division) indicating the involvement of UHMK1 in these processes at both post-translational and gene expression levels. Moreover, the GSEA analysis pointed to EMT as one of the enriched gene sets. Among the DEGs found in this study, *Prrx1* [40,41], *Pimreg* [42], *Sptbn1* [43,44], *Lox* [45], *Memo1* [46], and *Map3k3* [47,48] were previously implicated in EMT. Therefore, our study provides new evidence for the association of UHMK1 function with EMT.

In a previous report, we showed that the protein coded by *Pimreg* interacts with UHMK1 and is phosphorylated by this kinase *in vitro*, and suggested that the PIMREG-UHMK1 interaction could be important in controlling cell proliferation and gene expression [29]. Recently, we showed that PIMREG is involved in DNA damage response (DDR) [49]. Interestingly, our phosphoproteome analysis showed that UHMK1 does not only phosphorylate PIMREG, but also other proteins related to DDR, namely RAD54L, THRAP3, BCLAF1, and TPT1 [50–54], besides PIMREG. Phosphorylation on S128 of PIMREG was significantly upregulated in UHMK1 knockdown and downregulated in UHMK1^WT^ and UHMK1^K54R^ overexpression (although these phosphosites did not cross our fold change cutoff). The residue S128 in mouse corresponds to S131 in human, the reported phosphorylation site of UHMK1 [29]. Interestingly, as described here for SF1, phosphorylation of PIMREG by UHMK1 *in vivo*, exhibited a reverse phosphorylation pattern from what has been observed *in vitro* when using part of the protein or synthetic peptides as substrates. The fact that *Pimreg* was identified among the DEGs, points to a complex regulatory network between these proteins *in vivo*, triggering questions for further investigation.

UHMK1 regulated phosphosites of 30 proteins involved in cell cycle regulation. One third of the cell cycle-related proteins were also annotated as cytoskeleton-related, and the majority of them is also annotated in the BP terms “microtubule cytoskeleton organization”, “cell division” and “mitotic cell division”. To validate this finding, we performed *in vitro* functional assays to evaluate the phenotypes related to these cellular processes in UHMK1 overexpressing and depleted cells.

Our functional experiments demonstrated that UHMK1^WT^ overexpression decreased the number of colonies formed while UHMK1 knockdown decreased proliferation. Furthermore, UHMK1 knockdown reduced migration, while UHMK1 overexpression increased migration. These results are in line with previous reports, showing that modulation of UHMK1 affects cell migration, colony formation, and proliferation [12,15,24], and further support the data that UHMK1 regulates the phosphorylation of protein networks governing these cellular processes.

## Conclusion

A combination of phosphoproteome and transcriptome analysis allowed us to improve the understanding of the role of UHMK1 in cellular processes that have been previously associated with this kinase, particularly in RNA metabolism and splicing. We demonstrated for the first time the effect of UHMK1 modulation on alternative splicing events and on the phosphorylation of a variety of splicing factors, implicating UHMK1 as a novel splicing regulatory kinase. Our data also supports a function of UHMK1 in the modulation of transcription and gene expression, through the regulation of proteins that act in different layers controlling gene expression including chromatin modification, RNA pol II regulation, spliceosome machinery, and mRNA stability, besides rRNA cleavage and protein translation. The general picture of our study is that UHMK1 controls phosphorylation of proteins, gene expression, and alternative splicing of targets that are involved in key cellular processes such as cell cycle, cell division, and microtubule organization.

## Material and methods

### Cell lines and culture conditions

The murine NIH3T3 and the human HEK 293T cell lines were obtained from the *Deutsche Sammlung von Mikroorganismen und Zellkulturen* (DSMZ). Cells were cultured in Dulbecco’s modified Eagle’s medium (DMEM) supplemented with 10% of fetal bovine serum (FBS) and 1% of the antibiotic penicillin/streptomycin (PAN Biotech, Aidenbach, Germany), at 37 ºC and 5% CO_2_. The cell lines were tested for mycoplasma using MycoAlert™ PLUS Mycoplasma Detection Kit (Lonza, Basel, Switzerland).

### Plasmid construction

The wild type rat *Uhmk1* coding sequence (UHMK1^WT^), in frame with an N-terminal Flag peptide, was cloned into the pMSCV-IRES-YFP (MIY) retroviral vector. The UHMK1 kinase-dead mutant (UHMK1^K54R^) was generated by introducing the K54R mutation in the MIY-Flag-Uhmk1 construct, using the QuickChange II XL Site Directed Mutagenesis kit (Agilent Technologies, Santa Clara, CA, USA) and the following primers: forward 5’-CCCCGGCGCCCTCAGGCAGTTCCTG-3’, reverse 5’-CAGGAACTGCCTGAGGGCGCCGGGG-3’. For UHMK1 knockdown (UHMK1-KD), shRNA sequences targeting the murine *Uhmk1* gene were designed using the BLOCK-IT^™^ RNAi Designer tool (Invitrogen™), cloned into the MSCV-U3-H1-Stuffer entry vector digested with BglII and HindIII and subcloned into the retroviral vector pMSCV-puromycin-IRES-EGFP siRNA digested with the NotI and ScaI [55]. Target sequences are: shUHMK1#1: 5’-GCAAACAGTTCTGCTATTA-3’, shUHMK1#2: 5’-GCTGGATGATGATTACCTTGA-3’, shUHMK1#3: 5’-GCACTGGATGCTCTAATAA-3’. The scrambled control sequence is (shCTRL): 5’-GCATAGGCTCGAATTCTAA-3’.

### Generation of stable lines by retrovirus production and transduction of NIH3T3 cells

To produce retroviral particles carrying UHMK1^WT^, UHMK1^K54R^ or shRNA-UHMK1 sequences, HEK 293T cells were co-transfected with the retroviral construct and the packaging plasmid pCL-Eco [56], using Polyethylenimine (PEI). The virus-containing media (VCM), supplemented with protamine sulfate (5 µg/ml) was used for NIH3T3 cell infection. Transduced cells were sorted by Fluorescence-Activated Cell Sorting (FACS) based on YFP (UHMK1 overexpression) and GFP (UHMK1 knockdown) expression.

### Immunoblotting

Cells were lysed in lysis buffer [50 mM Tris pH 8.5, 150 mM NaCl and 1% Triton X-100 and protease inhibitor cocktail (Halt™ Protease Inhibitor Single-Use Cocktail; Thermo Fisher Scientific, Waltham, MA, USA)]. Total protein extracts were separated by electrophoresis on 10% SDS-polyacrylamide gel and transferred to a TransBlot® Turbo Mini Size polyvinylidene difluoride (PVDF) membrane using the TransBlot® Turbo system (BioRad Laboratories, Inc., Hercules, CA, USA). The membranes were blocked with low-fat milk and probed with primary antibody, followed by secondary antibodies conjugated to horseradish peroxidase (HRP). The proteins were detected by chemiluminescence with Pierce™ ECL Plus Western Blot Substrate (Thermo Fisher Scientific Scientific, Waltham, MA, USA) and images were captured with Fusion SL (Vilber Loumart, Eberhardzell, Germany). Primary antibodies were anti-UHMK1 (KIS-3B12, 1:10) [34], anti-GFP (sc-8334, 1:4000) and anti-Actin (sc-1616, 1:1000). Secondary antibodies were anti-rat (sc-2006, 1:2000) and anti-rabbit (sc-2313, 1:2000 or 1:8000). Except for anti-UHMK1, all antibodies were purchased from Santa Cruz Biotechnology, Dallas, TX, USA.

### Protein extraction, sample preparation and LC-MS/MS

UHMK1 overexpressing and depleted NIH3T3 cells were seeded in 100 mm plates at a density of 1.5×10^6^ cells/plate. After 24 h cells were harvested and lysed in lysis buffer [8 M urea, 80 mM Tris-HCL pH 7.6 and 1X Halt™ EDTA-free Protease Inhibitor Single-Use Cocktail (Thermo Fisher Scientific, Waltham, MA, USA) and 1X Phosphatase Inhibitor Cocktail 1, 2 and 3, (Sigma-Aldrich, St. Louis, MO, USA)] at constant rotation, for 45 min at 4°C. Whole-cell extracts were cleared by centrifugation and 300 µg of each protein extract was used for the subsequent steps. Reduction of disulfide bonds was performed using 10 mM DTT; cysteine residues were alkylated by incubating with 50 mM chloroacetamide; digestion was carried out using trypsin in a 1:50 enzyme-to-substrate ratio [57]. Samples were labeled with TMT6 reagent [58], the phosphopeptides were enriched using Fe-IMAC, and fractionation was performed using self-packed StageTips, as previously described [59]. Nanoflow LC-MS/MS was performed by coupling an UltiMate™ 3000 RSLCnano chromatography system to an Orbitrap Fusion Lumos Tribrid mass spectrometer (Thermo Fisher Scientific, Waltham, MA, USA). Samples were separated using a 90-min linear gradient from 4% to 32% LC solvent B (0.1% formic acid, 5% DMSO in acetonitrile) at a flow rate of 300 nl/min.

The mass spectrometer was operated in data-dependent acquisition mode. Full-scan MS spectra (m/z 360-1300) was recorded at a resolution of 60,000 and with an automatic gain control (AGC) target value of 4e5. Peptides were fragmented by collision-induced dissociation (CID) at 35% normalized collision energy with multistage activation in the ion trap and using and AGC target value of 5e4 and a maxIT of 60ms. Fragment ions were recorded in Orbitrap at a resolution of 30,000. For TMT quantification, peptides were fragmented in the ion trap as above with a maximum injection time (maxIT) of 120ms and AGC target value of 1.2e5. Fragment ions were subjected to synchronous precursor selection (SPS) and fragmented in the higher energy collisional dissociation (HCD) cell at a normalized collision energy of 55%. The resulting MS3 spectrum was recorded in the Orbitrap at a resolution of 50,000.

### Phosphoproteomics data analyses

The raw MS data were processed using MaxQuant v1.6.2.10 (https://www.maxquant.org) [60]. MS/MS spectra were searched against the UniProt mouse database supplemented with common contaminants. Carbamidomethylated cysteine, TMT-modified lysine side chain and peptide N-terminal were set as fixed modification, whereas oxidation (Met), N-terminal protein acetylation and phosphorylation (STY) were set as variable modifications. A target-decoy FDR of 1% was used at peptide spectrum match (PSM) level.

The MaxQuant results were further analyzed on the Perseus platform (https://maxquant.net/perseus/) [61]. Hits of the reverse and contaminant databases were removed. Relative abundances of phosphopeptides were determined by TMT reporter ion intensities from all PSMs matching to phosphopeptides. To normalize the quantification across different samples, TMT intensities of each TMT channel were summed up and normalized by median centering. The TMT intensities were then divided by the median intensity of all three replicates for each phosphopeptides. Ratios were further used for normalization. Only phosphopeptides found in all 3 replicates were considered for the analysis. Differentially phosphorylated phosphopeptides were defined by p-value < 0.05 and fold change of |1|. The identified phosphosites were searched in the databases PhosphositePlus (https://www.phosphosite.org/homeAction.action) [62] and PHOSIDA (http://141.61.102.18/phosida/index.aspx) [63].

### Gene Ontology and network analyses of the differentially phosphorylated proteins

Gene Ontology (GO) analysis and Reactome pathway analysis were carried out using the Protein Analysis Through Evolutionary Relationships -PANTHER (http://geneontology.org) [64]. Terms with FDR < 0.05 were considered statistically significant.

The differentially phosphorylated proteins and UHMK1 were searched for interactions using the STRING database (https://string-db.org) [65]. Interactions determined experimentally, registered in curated databases, extracted from textmining or co-expression data were considered, using a medium confidence (0.400) interaction score. Network was clustered using the MCL clustering algorithm, with an inflation parameter of 3.

### Consensus motif search and ULM motif analyses

Analysis of the consensus motif flanking the significantly regulated phosphosites was performed using pLogo [25]. We considered the phosphosites that are very likely direct substrates of UHMK1, *i*.*e*., all the phosphosites that were upregulated in UHMK1^WT^ overexpression, and downregulated in UHMK1-KD. Besides, we considered all phosphosites in UHMK1^K54R^. In total, 96 phospho-serines and 5 phospho-threonines were submitted to pLogo analysis. Search for ULM motif on the 28 RNA-related proteins (annotated in the RNA-related BP terms in the GO analysis) was performed using ScanProsite [66] and alignment in Clustal Omega [67]. We considered putative ULM domains those sequences that, beyond the conserved tryptophan, had at least 2 other conserved amino acids in the assigned positions (before or after the tryptophan), based on the known ULM motifs previously reported [26].

### RNA extraction, cDNA library preparation and RNA sequencing

An independent transduction of NIH3T3 cells overexpressing or depleted of UHMK1 was prepared for the RNA-seq experiment. 1×10^4^ cells of each sample were lysed in 100 µl of RLT Plus Lysis Buffer (Qiagen, Hilden, Germany) supplemented with 1% 2-mercaptoethanol. Replicates were harvested in 3 and 6 consecutive days, for UHMK1 knockdown and overexpression, respectively. A bulk RNA barcoding and sequencing protocol, named Prime-seq [32] was used for library preparation. Briefly, RNA was cleaned up using solid phase reversible immobilization (SPRI) beads (Sera Mag™, GE Healthcare Life Sciences, Chicago, IL, USA) in a homemade bead binding buffer containing 22% polyethylene glycol (PEG). cDNA was generated by Reverse Transcription mix containing the Maxima H Minus reverse transcriptase (Thermo Fisher Scientific, Waltham, MA, USA), oligo-dT primer E3V6NEXT and template switch primer E5V6NEXT. cDNA from all the samples was pooled together pre-amplified using KAPA HiFi HotStart polymerase (Roche, Basel, Switzerland) and SingV6 primer. Nextera libraries (5 replicates) were constructed from 0.8 ng of pre-amplified cleaned up cDNA using Nextera XT Kit (Illumina, Eindhoven, Netherlands). Index PCR was carried out using the custom P5 primer (P5NEXTPT5) and the i701 Nextera primer (IDT Technologies, Corallville, IA, USA). Libraries were size selected in a 2% E-Gel Agarose EX Gels (Life Technologies, Darmstadt, Germany), cut out in the range of 300–800 bp, and extracted using the MinElute Kit (Qiagen, Hilden, Germany), according to manufacturer’s recommendations.

Single-end sequencing was performed in the Illumina HiSeq 1500 system (Illumina, San Diego, CA, USA) with a coverage of 74 bp. Raw FASTQ data were processed with the pipeline zUMIs, version 0.2.0 [68] and mapped to the mouse genome mm10 using the software STAR [69]. Gene annotations were obtained from Ensembl (GRCm38.75).

### RNA-seq data analyses and target validation by qPCR

Differential expression between UHMK1-KD (shUHMK1#1, shUHMK1#2 and shUHMK1#3 data, combined and used as one pseudosample) and shCTRL was assessed using the edgeR/limma package [70,71]. Gene set enrichment analysis was performed using GSEA version 6.2 [72], with gene sets for *Mus musculus* obtained from GO2MSIG [73]. Results were regarded as significant with a p-value < 0.05.

RNA-seq validation was carried out in an independent transduction of NIH3T3 cells, by analyzing the expression of 8 differentially expressed genes (DEGs) by PCR array (qPCR). The DEGs chosen for validation were: *Cdca4, Fam198b, Pimreg, Lox, Map3k3, Oaz1, Prrx1* and *Sptbn1*. The cells were cultivated in the same conditions described for RNA-seq. Total RNA was extracted using RNeasy Mini Kit (Qiagen, Hilden, Germany). cDNA was produced using the iScript™ cDNA Synthesis Kit (Bio-Rad Laboratories, Hercules, CA, USA). qPCR was performed using PrimePCR custom plates (Bio-Rad Laboratories, Hercules, CA, USA), containing the lyophilized primers for the selected gene. The housekeeping genes *Gapdh* and *Hprt* were used as endogenous control and relative expression was calculated using the 2^-ΔΔCq^ method [74].

### Analysis of alternative splicing

For alternative splicing events (ASEs) detection in the mouse genome (mm10), we considered all raw sequencing reads from the replicates of each condition as four pseudosamples (UHMK1-KD, shCTRL, UHMK1^WT^ and EV) and identified ASEs using Bayesian inference followed by differential analysis from vast-tools pipeline (vast-tools diff module) [75,76] with |dPSI| ≥ 0.2 and MV|dPSI_at_95| ≥ 0.05 as significance thresholds. Additional non-default parameters for vast-tools diff module include: -S 1, -e 10, -m 0.05. ASEs were further categorized into functional classifications according to their predicted impact on the ORF: “neutral” for events that generate known functional isoforms or which do not alter the protein sequence (ex. an alternative exon); “protective” for events that reduce the occurrence of deleterious nucleotide sequences and therefore generate a functional protein (ex. removal of an intron/exon containing a premature stop codon) and “disruptive” which denote events that increase the frequency of disruptive sequences in the ORF (ex. inclusion of introns/exons containing premature stop codons or removal of essential exons for protein function) [77]. All post-processing data analysis and figure generation were conducted using custom python3.7 scripts, which are available upon request.

### Splicing reporter assays

HEK 293T cells were seeded in 24-well plates at a density of 9×10^4^ cells/well. After 24 h, the cells were co-transfected with 200 ng of the pTN24 reporter plasmid [78], MIY-UHMK1^WT^ and MIY-UHMK1^K54R^ (200 ng) construct alone or in combination with pcDNA-SF1[4] (300 ng of each construct). DNA was kept constant at 800 ng DNA in each well and 2 µl of Lipofectamine™ 2000 (Thermo Fisher Scientific, Waltham, MA, USA) was used as a transfection reagent. The cells were harvested 24 h later and assayed for luciferase and β-Galactosidase activity using Dual Light™ Luciferase and β-Galactosidase Reporter Gene Assay System (Thermo Fisher Scientific, Waltham, MA, USA). The measurements were performed in duplicates in an Infinite F200 Pro microplate reader (Tecan Group Ltd., Männedorf, Switzerland). Splicing was evaluated as a ratio of luciferase/ β-galactosidase. Parallel transfections were performed in the same conditions to assess gene expression of UHMK1^WT^, UHMK1^K54R^ and SF1 by qPCR.

### Colony forming assay

Cells were seeded in 6-well plates at a density of 750 cells/well. After 6 days of incubation under normal culturing conditions, the colonies were fixed with 70 % ethanol for 10 min, stained with 0.05% crystal violet for 10 min, and washed with water. The colonies were counted using an inverted microscope (Motic, Hong Kong, China). A colony was defined by a minimum of 50 cells.

### Proliferation assay

Proliferation was evaluated by BrdU incorporation. 2×10^5^ cells were seeded in 6-well plates. After 24 h, 10 µM of BrdU was added and incubated for 2 h. BrdU staining was carried out with the APC BrdU Flow Kit (BD Biosciences, Franklin Lakes, NJ, USA). Flow cytometry analyses were performed using a FACS Canto (BD Biosciences, Franklin Lakes, NJ, USA).

### Cell viability assay

Cells were seeded in 96-well plates at a density of 8×10^3^ cells/well. After 24 h, 20 µl of Cell Titer Blue® reagent (Promega, Madison, WI, USA) was added to the cells and incubated for 4 h at 37 ºC, protected from light. Fluorescence was measured using Promega GloMax® Microplate Reader, excitation filter: 520 nm, emission: 580-640.

### Apoptosis assay

Cells were seeded in 12-well plates at a density of 1×10^5^ cells/well. After 24h, cells were washed with ice-cold DPBS plus 5 mM EDTA and stained with annexin V and PI (BD Biosciences, Franklin Lakes, NJ, USA) for 15 min at RT. Analysis was carried out using FACS Canto (BD Biosciences, Franklin Lakes, NJ, USA).

### Migration assay

Cell cultures were depleted from serum (1% FBS-containing medium without antibiotics) for 16 -20 h. After that period, 1×10^5^ cells were seeded directly over an 8 µm membrane in a 96-well Boyden chamber plate. The lower compartment of the chamber was filled with 10% FBS-containing medium. After 24h, the cells that migrated through the membrane were fixed with 70% ethanol and stained with 0.05% crystal violet, as previously described [79]. Images were acquired using Microscope Leica DMi8 at 10x magnification.

### Statistical analyses

Statistical analyses (of splicing reporter assays, RNA-seq validation, and functional assays) were carried out on GraphPad Prism version 7.0 for MacOS X (GraphPad Software, San Diego, CA, USA, available at https://www.graphpad.com). The function Column Statistics was used to evaluate parameters such as median, mean, coefficient of variation, kurtosis, skewness, and normality (Shapiro-Wilk test). Based on these parameters, the tests chosen to evaluate 3 groups or more were One-way ANOVA followed by Bonferroni’s multiple comparison test or Kruskal-Wallis test followed by Dunn’s multiple comparison test. For 2 groups, unpaired Student T test or Mann-Whitney test were used. A confidence interval (CI) of 0.95 was set and therefore p-values lower than 0.05 were regarded as statistically significant.

## Data availability

The RNA-seq data from UHMK1-overexpressing or -depleted NIH3T3 cells are available in the Gene Expression Omnibus (GEO) repository, accession number GSE199768. The UHMK1 mass spectrometry proteomics data have been deposited to the ProteomeXchange Consortium via the Proteomics Identification Database (PRIDE) partner repository with the dataset identifier PXD033353.

## CRediT authorship contribution statement

**Vanessa C. Arfelli:** Investigation, Formal analysis, Visualization, Validation, Writing - Original Draft. **Yun-Chien Chang:** Investigation, Formal analysis. **Johannes W. Bagnoli:** Investigation, Formal analysis. **Paul Kerbs:** Formal analysis, Visualization. **Felipe E. Ciamponi:** Formal analysis, Visualization. **Laissa M. S. Paz:** Investigation. **Katlin B. Massirer:** Formal analysis, Supervision. **Wolfgang Enard:** Supervision, Resources. **Bernhard Küster:** Supervision, Resources. **Philipp A. Greif:** Conceptualization, Resources, Supervision, Writing - Review & Editing. **Leticia Fröhlich Archangelo:** Conceptualization, Supervision, Writing - Original Draft, - Review & Editing, Project administration, Funding acquisition.

## Acknowledgements

We thank Dr. Ian C. Eperon for the pTN24 reporter plasmid, Dr. Alexandre Maucuer for the pcDNA-SF1 construct, Dr. Saghi Ghaffari for the MSCV-U3-H1-Stuffer and pMSCV-puromycin IRES2-EGFP siRNA vectors, and Dr. Alexander Faussner for the support in the splicing reporter assay readouts. This research was supported by the Coordenação de Aperfeiçoamento Pessoal de Nível Superior (CAPES/PDSE, grant to VCA and to FEC), German Science foundation (DFG-SFB1309, grant to YCC) and the Fundação de Amparo à Pesquisa de São Paulo (FAPESP 2020/02006-0 to KBM and 2014/01458-3 to LFA). PAG acknowledges support by the Wilhelm Sander-Stiftung (Förderantrag Nr. 2014.162.3) and the Munich Clinician Scientist Program (MCSP).

