## Supplemental Figures for "UHMK1 is a novel splicing regulatory kinase"

### Supplementary Figures

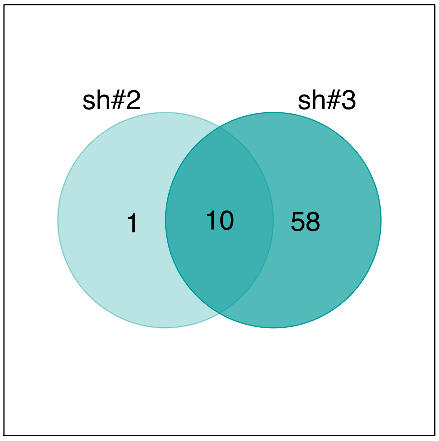


**Sup. Figure 1.** Comparison between the number of differentially phosphorylated proteins (DPPs) in UHMK1 knockdown cells driven by shUHMK1#2 (sh#2) and shUHMK1#3 (sh#3) sequences.

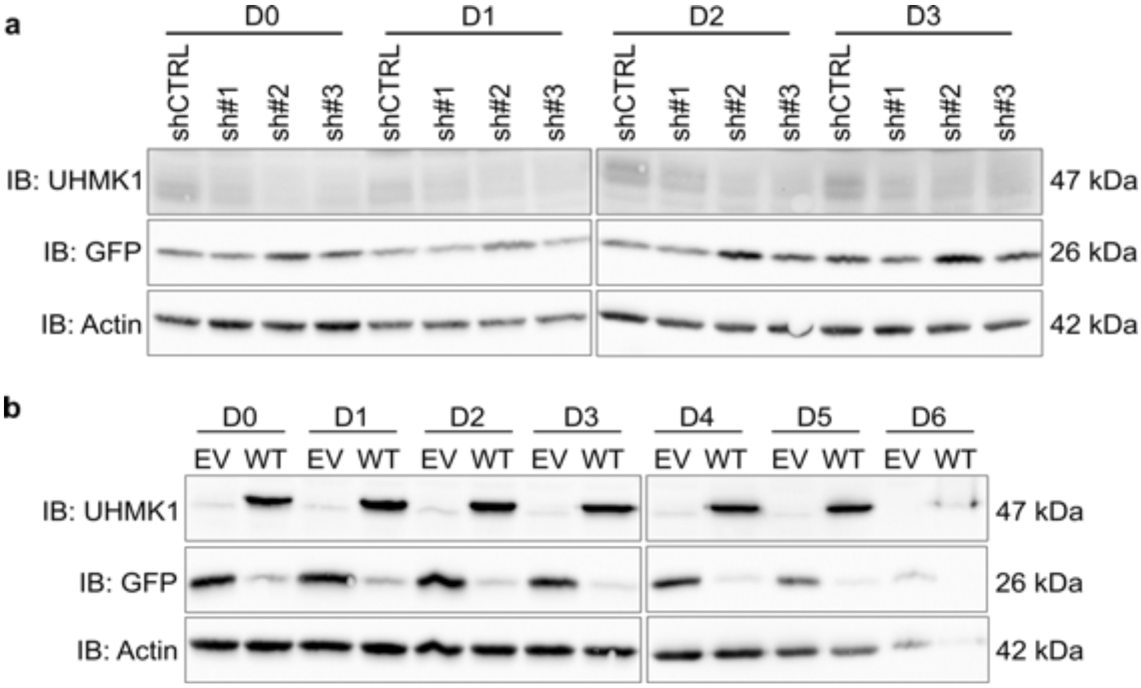


**Sup. Figure 2.** Independent transduction of NIH3T3 cells for the RNA-seq experiment. Western blot confirming efficient (a) UHMK1 knockdown mediated by shUHMK1#1 (sh#1), shUHMK1#2 (sh#2), and shUHMK1#3 (sh#3) compared to shCTRL cells, and (b) overexpression of UHMK1$^{WT}$ (WT) compared to the empty vector (EV) control. D0 – D6 refer to the consecutive days in which the cells were harvested for RNA extraction.

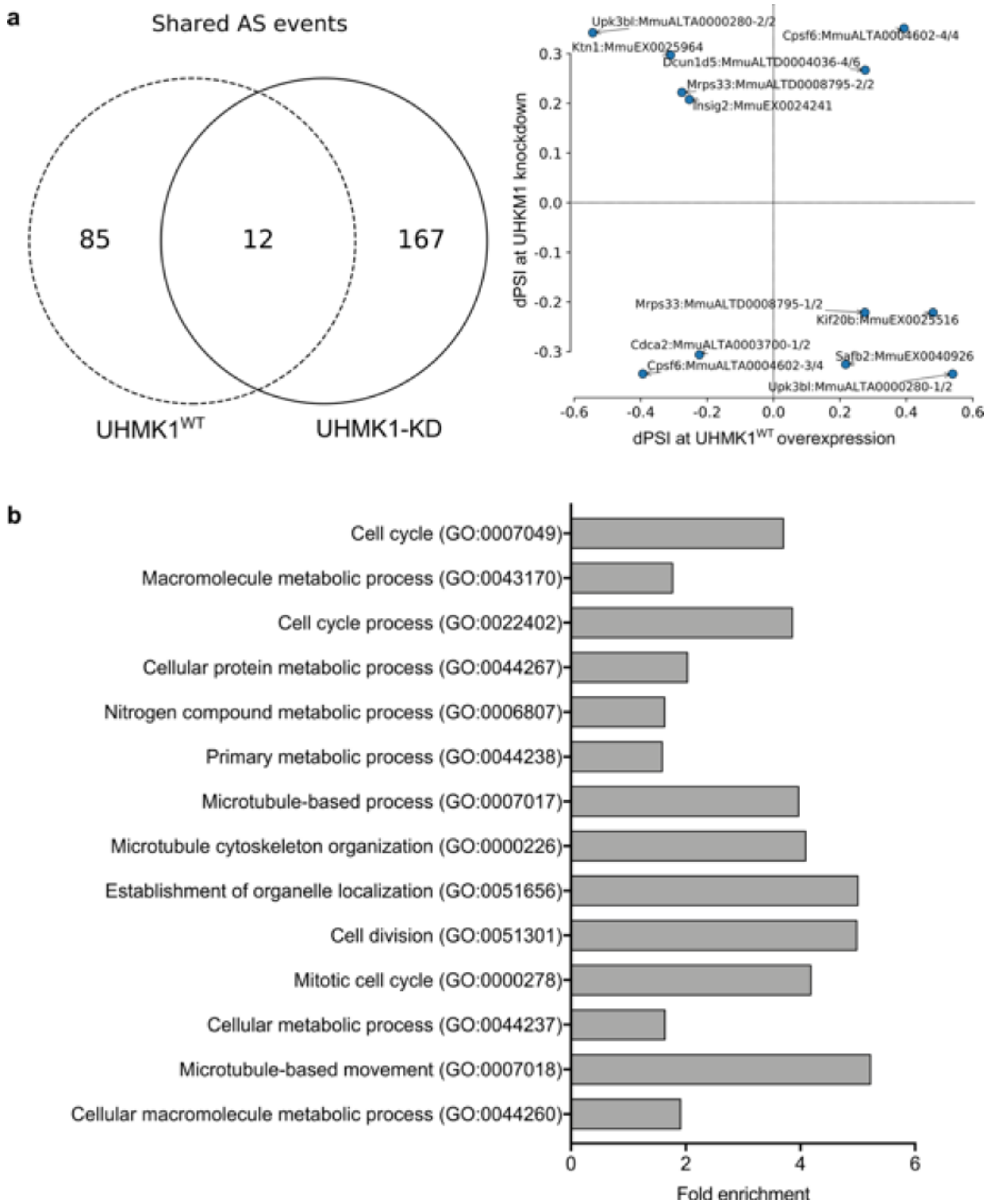


**Sup. Figure 3.** UHMK1 impact on splicing. (a) Shared alternative splicing events (ASEs) between UHMK1$^{WT}$ overexpression and UHMK1 knockdown (UHMK1-KD). (b) Biological Process terms returned from Gene Ontology analysis performed with the target genes involved in ASEs in UHMK1 knockdown cells (FDR < 0.05). No significant results were retrieved with the target genes involved in ASEs in UHMK1$^{WT}$ overexpressing cells.

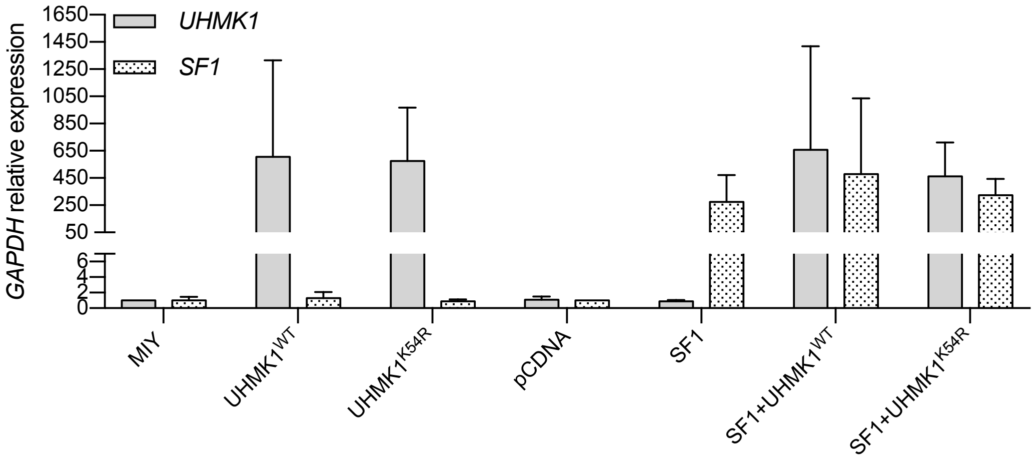


**Sup. Figure 4.** Verification by qPCR of *UHMK1* and *SF1* expression levels relative to *GAPDH* in parallel transfections, performed with the same experimental setting of the pTN24 splicing reporter assay.

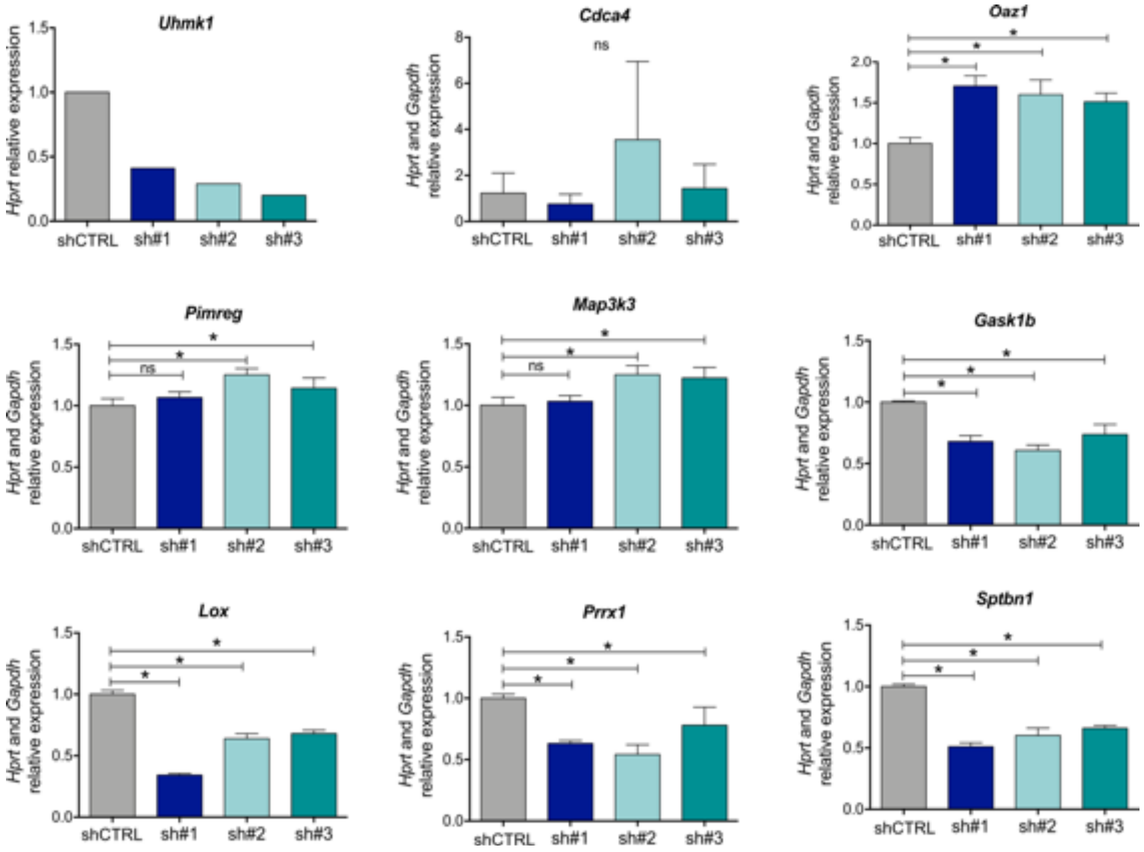


**Sup. Figure 5.** UHMK1 knockdown effect in gene expression was validated on independent samples. Expression of *Uhmk1* and the 8 selected genes relative *Hrpt* and *Gapdh* expression. PCR array for validation was performed twice in duplicates. Error bars represent standard deviation from 4 measurements. Asterisks represent significant difference (Mann-Whitney test, $p < 0.05$) of UHMK1 knockdown cells mediated by shUHMK1#1 (sh#1), shUHMK1#2 (sh#2), and shUHMK1#3 (sh#3) relative to control cells (shCTRL).

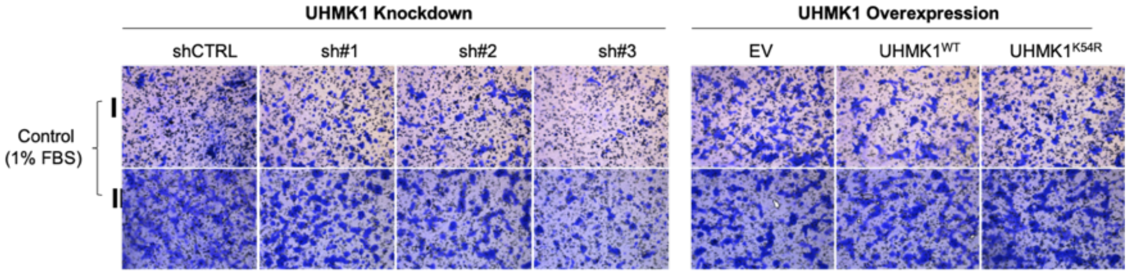


**Sup. Figure 6. Effect of UHMK1 in migration.** Control panels showing spontaneous migration towards the lower chamber containing 1% FBS. Images were taken in the center of the well and can be compared to the imagens presented in Figure 8e. 10 x magnification, Microscope Leica DMi8.

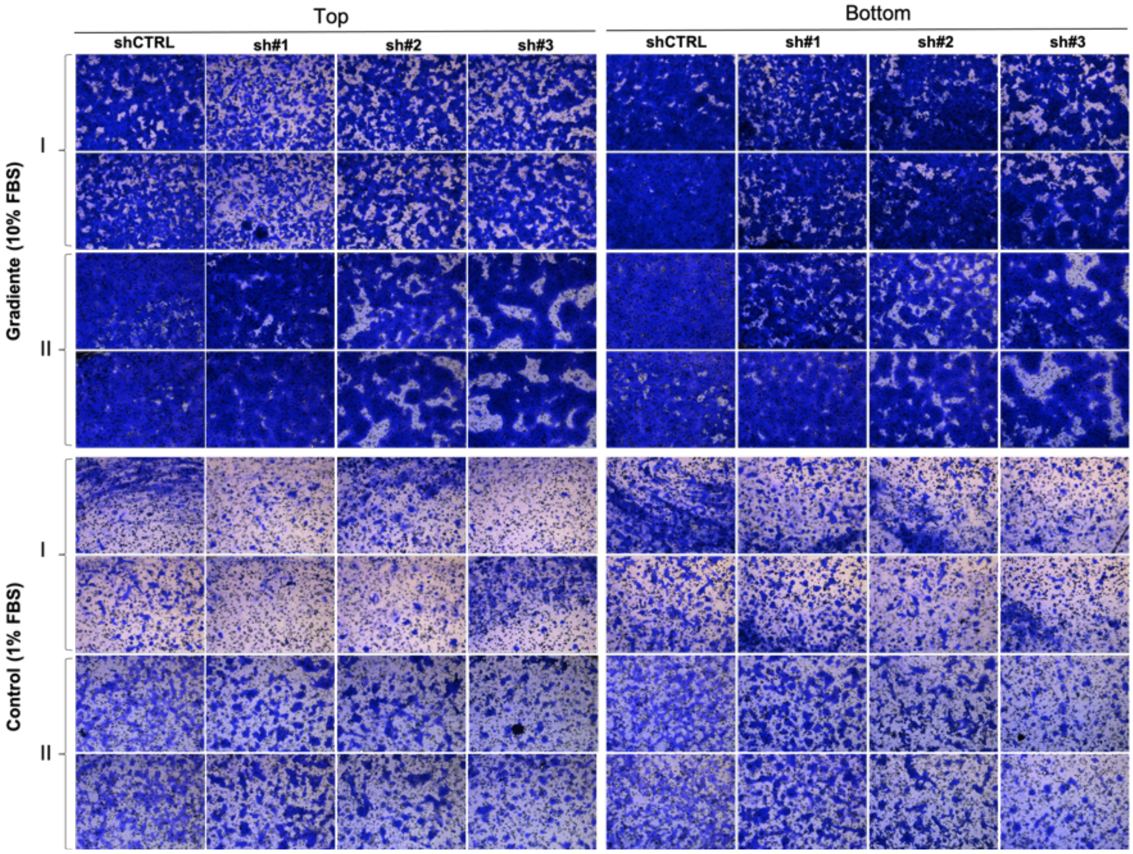


**Sup. Figure 7. Effect of UHMK1 knockdown in migration.** NIH3T3 cells depleted of UHMK1 migrated less through an 8 μM-pore membrane, in comparison to shCTRL (upper panel, gradient 10% FBS). Images from two independent experiments (I and II), each in duplicate, acquired on the top and bottom of the well, relative to the orientation of the circumference in the plate. The Control panel represents spontaneous migration, as the lower chamber had 1% FBS, the same FBS concentration present in the upper chamber with the cells. 10 x magnification, Microscope Leica DMi8.

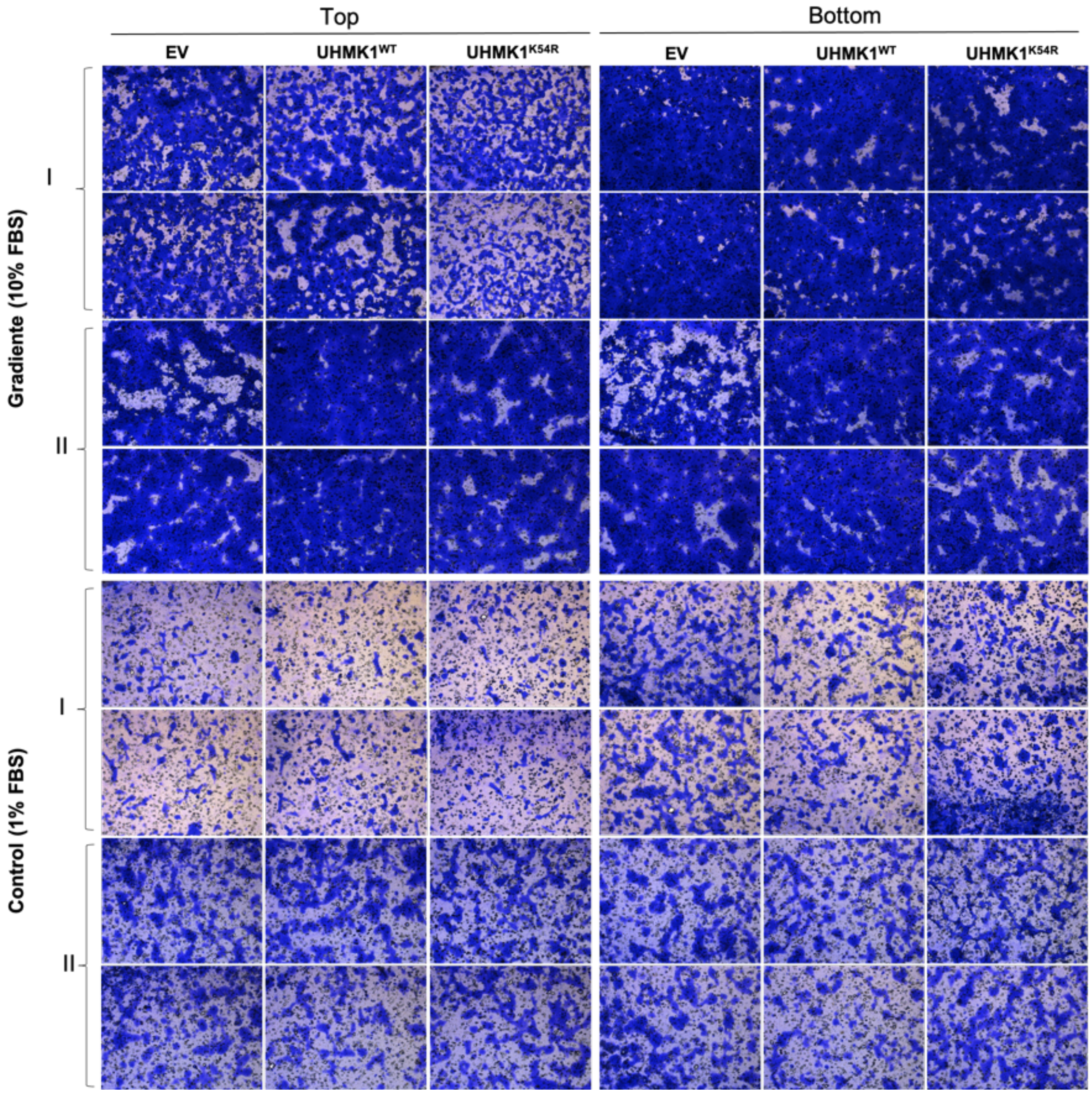


**Sup. Figure 8. Effect of UHMK1 overexpression in migration**. Images from two independent experiments (I and II), each in duplicate, acquired on the top and bottom of the well, relative to the orientation of the circumference in the plate. NIH3T3 cells overexpressing UHMK1$^{WT}$ seemed to migrate more through an 8 μM-pore membrane (except in Top and Bottom of experiment I), in comparison to the empty vector (EV) control (upper panel, gradient 10% FBS). In these exact same conditions, UHMK1$^{K54R}$ overexpression migrated less. The Control panel represents spontaneous migration, as the lower chamber had 1% FBS, the same FBS concentration present in the upper chamber with the cells. 10 x magnification, Microscope Leica DMi8.
