## Supplemental Tables for "UHMK1 is a novel splicing regulatory kinase"

### Supplementary Tables

Sup. Table 1 - The 117 DPPs, phosphopeptides and phosphosites

| Gene names | Protein code (Uniprot) | Phosphopeptides | (STY) sites | Phospho-sites | log2 fold change | | | |
|---|---|---|---|---|---|---|---|---|
| | | | | | shUHMK1 #2 | shUHMK1 #3 | UHMK1$^{WT}$ | UHMK1$^{K54R}$ |
| **Proteins differentially phosphorylated in all conditions** | | | | | | | | |
| ‡Avil | O88398 | _pYpYPVEVLLK_ | Y;Y | 758; ‡759 | -2.45455 | -2.22162 | 1.38213 | 1.23678 |
| ‡Gm7995 | K7N665 | _QpSISDSIDK_ | S | 80 | -2.08122 | -1.96843 | 1.54604 | 1.29922 |
| Lmna - I | P48678 | _ASpSHSSQSQGGGSVTK_ | S | 404 | | | 1.98789 | 1.48258 |
| Lmna - II | P48678 | _ASpSHSpSQSQGGGSVTK_ | S;S | 404;407 | | 1.03270 | 1.05241 | |
| ‡Rad54l | P70270 | _pTLQCIpTLMWTLLR_ | T;T | 190;195 | 4.76120 | -1.07729 | 4.37880 | 4.62590 |
| Sf1 - I | D3YVH4 | _SPpSPEPIYNSEGKR_ | S | 56 | 3.08121 | 2.63871 | | |
| Sf1 - II | D3YVH4 | _SPpSPEPIYNSEGK_ | S | 56 | 2.91822 | 1.98151 | -2.44725 | |
| Sf1 - III | D3YVH4 | _TGDLGIPPNPEDRpSPpSPEPIYNSEGKR_ | S;S | 54;56 | | | | -1.19762 |
| ‡Stxbp4 | Q9WV89 | _AQLADYpSDQNK_ | S | 396 | -2.04610 | -2.05198 | 1.36103 | 1.21770 |
| Tppp | Q7TQD2 | _ERFDQpSGK_ | S | 182 | | 4.93442 | 4.12719 | 4.76845 |
| **Proteins differentially phosphorylated in UHMK1-KD and UHMK1$^{K54R}$** | | | | | | | | |
| Atp2b1 - I | A0A1W2P772 | _IEDpSEPHIPLIDDTDAEDDAPTKR_ | S | 252 | | -1.62882 | | |
| Atp2b1 - II | A0A1W2P772 | _IEDpSEPHIPLIDDTDAEDDAPTK_ | S | 252 | | -1.55312 | | 1.04663 |
| Atp2b1 - III | A0A1W2P772 | _NSpSPPPpSPNK_ | S | 275;279 | | -1.34461 | | |
| Canx - I | P35564 | _AEEDEILNRpSPR_ | S | 582 | | -1.51576 | | 1.14060 |
| Canx - II | P35564 | _QKpSDAEEDGVTGSQDEEDSKPK_ | S | 553 | | -1.40307 | | 1.17531 |
| Canx - III | P35564 | _QKpSDAEEDGVTGpSQDEEDSKPK_ | S;S | 553;563 | | -1.18887 | | 1.30555 |
| Cd44 | Q3U8S1 | _pSQEMVHLVNKEPSETPDQCMTADETR_ | S | 329 | | -1.02502 | | 1.38557 |
| Eif4g1 - I | E9PVC6 | _AApSLTEDRGR_ | S | 1164 | | 1.06497 | | |

Sup. Table 1 - The 117 DPPs, phosphopeptides and phosphosites

| Gene names | Protein code (Uniprot) | Phosphopeptides | (STY) sites | Phospho-sites | log2 fold change | | | |
|---|---|---|---|---|---|---|---|---|
| | | | | | shUHMK1 #2 | shUHMK1 #3 | UHMK1$^{WT}$ | UHMK1$^{K54R}$ |
| Eif4g1 - II | E9PVC6 | _pSSLpSRERGEK_ | S;S | 1099;1102 | | | | -1.07514 |
| Pla2g4a | P47713 | _HIVSNDSSDpSDDEAQGPKGTENEEAEK_ | S | 437 | | 2.31953 | | -1.04078 |
| Sec62 - I | Q8BU14 | _EETPGpTPK_ | T | 158 | | -1.57116 | | 1.31984 |
| Sec62 - II | Q8BU14 | _KEETPGpTPK_ | T | 158 | | -1.27313 | | 1.03731 |
| Slain2 - I | Q8CI08 | _NpSPRPpSPK_ | S;S | 350;354 | | -1.04298 | | |
| Slain2 - II | Q8CI08 | _GTFpSDQELDAQpSLDDEDDSLQHAVHPALNR_ | S;S | 316;324 | | | | 1.05420 |
| Srrm2 - I | Q8BTI8 | _AGRpSRpSPATK_ | S;S | 482;484 | | 2.07805 | | |
| Srrm2 - II | Q8BTI8 | _ERSGAGpSPPGKR_ | S | 1179 | | 1.34044 | | |
| Srrm2 - III | Q8BTI8 | _NHpSGSRTPPVALSSSR_ | S | 2052 | | 1.47945 | | |
| Srrm2 - IV | Q8BTI8 | _PApSPKKPPPGER_ | S | 2618 | | | | -1.70386 |
| Srrm2 - V | Q8BTI8 | _SKpTPPRQSR_ | T | 805 | | | | -1.22822 |
| Tmem245 | D3YWD3 | _GGPAEAPpSPR_ | S | 12 | | -1.51796 | | 1.03213 |
| **Proteins differentially phosphorylated in UHMK1$^{WT}$ and UHMK1$^{K54R}$** | | | | | | | | |
| Bop1 | P97452 | _AEETpSEELAQAAPLCSR_ | S | 84 | | | 1.81487 | 2.08761 |
| Fosl2 | P47930 | _GSSGpSPAHAESYSSGGGGQQK_ | S | 19 | | | 1.30301 | 1.07183 |
| Ftsj3 | Q9DBE9 | _ALDISLpSpSEEEEEGDEEEAVAETK_ | S;S | 335;336 | | | 1.40309 | 1.66438 |
| Iqgap1 | Q9JKF1 | _(ac)pSAAEEVDGLGVVRPHYGSVLDNER_ | S | 2 | | | -1.18835 | -1.39366 |
| Pde8a | O88502 | _HpSpSLARIHpSMMIEAPITK_ | S | 375; 376; 382 | | | -1.98523 | -1.66279 |
| Snrnp200 | Q6P4T2 | _EEApSDDDMEGDEAVVR_ | S | 225 | | | 1.50799 | 1.68938 |
| ‡Tbcd | Q8BYA0 | _VLSNEPAApSAAEEEVEDDALVR_ | S | 10 | | | -1.02385 | -1.12887 |
| **Proteins differentially phosphorylated in UHMK1$^{WT}$ and UHMK1-KD** | | | | | | | | |
| Med19 | Q8C1S0 | _NRHpSPDHPGMGSSQASSSSSLR_ | S | 226 | | 1.30193 | 1.86052 | |

Sup. Table 1 - The 117 DPPs, phosphopeptides and phosphosites

| Gene names | Protein code (Uniprot) | Phosphopeptides | (STY) sites | Phospho-sites | log2 fold change | | | |
|---|---|---|---|---|---|---|---|---|
| | | | | | shUHMK1 #2 | shUHMK1 #3 | UHMK1$^{WT}$ | UHMK1$^{K54R}$ |
| **Proteins differentially phosphorylated solely in UHMK1-KD** | | | | | | | | |
| Abcf1 | Q6P542 | _GGNVFEALIQDDpSEEEEEEEENRVLK_ | S | 138 | | 1.25963 | | |
| Ahnak | E9Q616 | _LQGpSGVSLASKK_ | S | 5617 | | 1.09783 | | |
| Apobr | Q8VBT6 | _NpSWATEPTLVLDTEAK_ | S | 398 | | -1.24560 | | |
| ‡Arid1a | E9QAQ7 | _IELLPpSR_ | S | 1870 | | -1.07627 | | |
| Atp2a2 | O55143 | _EFDELSPpSAQR_ | S | 663 | | -1.19145 | | |
| C4orf3 homolog | Q99M08 | _RGpSFEAGR_ | S | 19 | | -1.14338 | | |
| C7orf50 homolog | Q9CXL3 | _ETASTLVQEASPELpSPEERR_ | S | 52 | 1.15365 | 1.27778 | | |
| Ccny | Q8BGU5 | _SApSADNLILPR_ | S | 326 | | -1.01253 | | |
| Ccnyl1 | E9Q226 | _SLpSADNFIGIQR_ | S | 276 | | -1.22030 | | |
| Cers2 | Q924Z4 | _LIEDERpSDREEpTEpSpSEGEETAAGAGAK_ | S;T | 341;348; 349;346 | | -1.04815 | | |
| Chfr | Q810L3 | _SSLVANGELSSLpSPVFQDK_ | S | 231 | | -1.11961 | | |
| Eef2 - I | P58252 | _FpTDTRKDEQER_ | T | 57 | | 1.40100 | | |
| Eef2 - II | P58252 | _AGETRFpTDTR_ | T | 57 | | 1.19379 | | |
| ‡Efr3a | A0A1D5RLL3 | _LTVPYVPQVTDEDRLpSR_ | S | 719 | | -1.00414 | | |
| Eif4ebp1 | Q60876 | _FLMECRNpSPVAKpTPPK_ | S;T | 64;69 | | 1.14876 | | |
| Hacd3 | Q8K2C9 | _WLDEpSDAEMELR_ | S | 114 | | -1.08912 | | |
| Irf2bp2 | E9Q1P8 | _RPApSVSSAAAEHEAREPSK_ | S | 226 | | 1.11709 | | |
| Luc7l | Q9CYI4 | _TASRRpSEEK_ | S | 363 | | 1.06374 | | |
| Map4 | P27546 | _PTLLANGDHGMEGNNTAGpSPTDFLEER_ | S | 99 | | -1.04821 | | |
| Mapre2 | E9Q6X0 | _SHHANpSPTAGAAK_ | S | 157 | | 1.00670 | | |
| Mcm3 | P25206 | _DGESYDPYDFSEAETQMPQVHpTPKTDDSQEK_ | T | 719 | | 1.16741 | | |

Sup. Table 1 - The 117 DPPs, phosphopeptides and phosphosites

| Gene names | Protein code (Uniprot) | Phosphopeptides | (STY) sites | Phospho-sites | log2 fold change | | | |
|---|---|---|---|---|---|---|---|---|
| | | | | | shUHMK1 #2 | shUHMK1 #3 | UHMK1$^{WT}$ | UHMK1$^{K54R}$ |
| Mtdh | Q80WJ7 | _SETNWEpSPK_ | S | 565 | | -1.41913 | | |
| Ncor2 | F8VQL9 | _pTPELPLAPR_ | T | 1441 | | -1.13210 | | |
| Npm1 | Q61937 | _pTPKpTPKGPSSVEDIK_ | T; T | 232;235 | 1.10225 | 1.25797 | | |
| Nucks1 | Q80XU3 | _VVDYSQFQEpSDDADEDYGRDSGPPAK_ | S | 19 | | 1.07093 | | |
| Palm | Q9Z0P4 | _EPAPLNGpSAAELPATK_ | S | 345 | | -1.26365 | | |
| Pgrmc1 | O55022 | _EGEEPTVYpSDDEEPKDETAR_ | S | 181 | | -1.05201 | | |
| Piezo1 | F6PYU5 | _TApSELLLDR_ | S | 996 | | -1.05689 | | |
| Ppfibp1 | Q8C8U0 | _TApSAPNLAETEKETAEHLNLAGTSR_ | S | 500 | | -1.40183 | | |
| Ppig | A2AR02 | _MRVSpSGER_ | S | 374 | | 1.32091 | | |
| Prpf38a | Q4FK66 | _LERVPpSPDHR_ | S | 209 | | 1.30595 | | |
| Prpf4b | Q61136 | _pSRpSLERK_ | S;S | 394;396 | | 1.00554 | | |
| Ptpn21 | A0A1Y7VNU2 | _RPVVGAVpSVPELTNVQLQAQDYPAPNIMR_ | S | 538 | | 1.09481 | | |
| Safb;Safb2 | S4R1M2;Q80YR5 | _VVTNARpSPGAR_ | S | 465 | | 1.02036 | | |
| Scarf2 | P59222 | _NEAGGLSLpSPpSPER_ | S;S | 638;640 | | -1.59002 | | |
| Scrib - I | Q80U72 | _LpSPDFVEELR_ | S | 1548 | | -1.22448 | | |
| Scrib - II | Q80U72 | _NpSLESISSIDR_ | S | 1206 | | -1.04444 | | |
| Sec61b | Q9CQS8 | _PGPTPSGTNVGSSGRpSPSK_ | S | 17 | | -1.37402 | | |
| Slc16a1 - I | P53986 | _LKpSKEpSLQEAGK_ | S | 210;213 | | -1.30723 | | |
| Slc16a1 - II | P53986 | _SKEpSLQEAGK_ | S | 213 | | -1.28001 | | |
| Slc1a3 | P56564 | _DVEMGNpSVIEENEMK_ | S | 512 | | -1.82517 | | |
| Slc33a1 | Q99J27 | _RDpSVGGEGDREVLLGDAGPGDLPK_ | S | 42 | | -1.02401 | | |
| Slc4a7 - I | F8VQC9 | _GpSLLQIPVK_ | S | 1057 | | -2.08344 | | |

Sup. Table 1 - The 117 DPPs, phosphopeptides and phosphosites

| Gene names | Protein code (Uniprot) | Phosphopeptides | (STY) sites | Phospho-sites | log2 fold change | | | |
|---|---|---|---|---|---|---|---|---|
| | | | | | shUHMK1 #2 | shUHMK1 #3 | UHMK1$^{WT}$ | UHMK1$^{K54R}$ |
| Slc4a7 - II | F8VQC9 | _KHpSDPHLLER_ | S | 247 | | -1.04070 | | |
| Slc4a7 - III | F8VQC9 | _GNGSGGpSRENSTVDFSK_ | S | 284 | | -1.31699 | | |
| Slc4a7 - IV | F8VQC9 | _NGILASPQpSAPGNLDNSK_ | S | 263 | | -1.39861 | | |
| Slc4a7- V | F8VQC9 | _pSFADIGK_ | S | 238 | | -1.44249 | | |
| Slc4a7 - VI | F8VQC9 | _MLQDDEDTVHLPFERGpSLLQIPVK_ | S | 1057 | | -1.53629 | | |
| Snap23 | B0R030 | _ATWGDGGDNpSPSNVVSK_ | S | 110 | | -1.13705 | | |
| Son | H9KV15 | _GRRpSVpSK_ | S | 1910;1912 | | 1.36006 | | |
| Sptbn1 | Q62261 | _TpSpSKESpSPVPpSPTLDR_ | S;S;<br>S;S | 2159;216;<br>2164;216; | | -1.13851 | | |
| Srsf2 | Q62093 | _VDNLTYRTpSPDTLRR_ | S | 26 | | 1.22582 | | |
| Srsf4 | Q542V3 | _SHpSPSRHDpSK_ | S;S | 291;297 | | 1.18710 | | |
| Stmn1 - I | P54227 | _ApSGQAFELILpSPR_ | S;S | 16;25 | | -1.14144 | | |
| Stmn1 - II | P54227 | _ApSGQAFELILSPR_ | S | 16 | | -1.24510 | | |
| Stx4a;Stx4 | D6RJ29 | _QGDNIpSDDEDEVR_ | S | 15 | | -1.02267 | | |
| ‡Tnrc6b | A0A2I3BRG1 | _DDEPSGWEEPpSPQSISR_ | S | 114 | -1.78131 | -1.70440 | | |
| Tpd52l2 | Q3TUJ9 | _pSFEDRVGTIK_ | S | 146 | 1.85874 | 2.33890 | | |
| Usp39 | Q3TIX9 | _EPEAASSRGpSPVR_ | S | 46 | 1.00392 | 1.24291 | | |
| Vps4a | Q8VEJ9 | _GSDpSDSEGDNPEK_ | S | 97 | | -1.06183 | | |
| Zdhhc5 | Q8VDZ4 | _GDpSLKEPTSIADSSR_ | S | 380 | | -1.45187 | | |
| **Proteins differentially phosphorylated solely in UHMK1$^{WT}$** | | | | | | | | |
| Abi1 | B7ZCU4 | _LGSQHpSPGR_ | S | 225 | | | 1.07337 | |
| Add1 | Q9QYC0 | _YSDVEVPASVTGHSFASDGDSGTCpSPLR_ | S | 431 | | | 1.37010 | |
| Ankrd17 | E9QKG6 | _EHYPVSSPSpSPSPPAQPGGVSR_ | S | 1790 | | | -1.24956 | |
| Arfgap2 | Q99K28 | _HGTDLWIDSMNSAPSHpSPEKK_ | S | 145 | | | 1.08437 | |

Sup. Table 1 - The 117 DPPs, phosphopeptides and phosphosites

| Gene names | Protein code (Uniprot) | Phosphopeptides | (STY) sites | Phospho-sites | log2 fold change | | | |
|---|---|---|---|---|---|---|---|---|
| | | | | | **shUHMK1 #2** | **shUHMK1 #3** | **UHMK1$^{WT}$** | **UHMK1$^{K54R}$** |
| Bclaf1 | Q8K019 | _NTPSQHSHSIQHpSPER_ | S | 267 | | | 1.40929 | |
| Cobll1 | Q3UMF0 | _DPQLpSPEQHPSSLSER_ | S | 993 | | | 1.21068 | |
| Cul4b | A2A432 | _DSApSPSTSSFCLGVPVATSSHVPIQK_ | S | 149 | | | 1.08308 | |
| Dnajc5 | A2AUE1 | _SLpSTSGESLYHVLGLDK_ | S | 10 | | | 1.33885 | |
| Edc3 | Q8K2D3 | _HNpSWSSSSR_ | S | 161 | | | 1.45147 | |
| Eef1g | Q9D8N0 | _VLSAPPHFHFGQTNRpTPEFLR_ | T | 46 | | | 1.23006 | |
| Hnrnpm | Q9D0E1 | _GCGVVKFEpSPEVAER_ | S | 700 | | | 1.52853 | |
| Igf2r - I | Q07113 | _AEALSSLHGDDQDpSEDEVLTVPEVK_ | S | 2401 | | | 1.09894 | |
| Igf2r - II | Q07113 | _LVSFHDDpSDEDLLHI_ | S | 2476 | | | 1.15429 | |
| Irf2bpl - I | Q8K3X4 | _RNpSSpSPVSPASVPGQR_ | S;S | 636;638 | | | -1.35291 | |
| Irf2bpl - II | Q8K3X4 | _RNpSSSPVpSPASVPGQR_ | S;S | 636;641 | | | -1.08179 | |
| L3mbtl2 | Q80XB0 | _EAGELPTpSPLHLFSSANNR_ | S | 67 | | | 1.23812 | |
| Mcm6 | Q3ULG5 | _HVDEFpSPR_ | S | 413 | | | 1.37883 | |
| Mphosph10 - I | Q810V0 | _VTFALPDDEAEDTpSPLAVK_ | S | 346 | | | 1.75154 | |
| Mphosph10 - II | Q810V0 | _VTFALPDDEAEDTpSPLAVKQESDEVK_ | S | 346 | | | 1.37964 | |
| Numa1 | E9Q7G0 | _VSSETHQGPGpTPESK_ | T | 1982 | | | 1.15276 | |
| Pdha1 - I | P35486 | _YHGHSMpSDPGVSYR_ | S | 295 | | | 1.08250 | |
| Pdha1 - II | P35486 | _YHGHpSMSDPGVSYR_ | S | 293 | | | 1.09127 | |
| Pik3c3 | E9Q824 | _DGDESpSPILTSFELVK_ | S | 244 | | | 1.57159 | |
| Ppp4r2 | Q0VGB7 | _GHSDSSASESEVSLLpSPVK_ | S | 226 | | | 1.17800 | |
| Prrc2b - I | F8WHT3 | _SPDEALPGGLGSHpSPYALER_ | S | 1493 | | | 1.58497 | |
| Prrc2b - II | F8WHT3 | _pSPDEALPGGLGSHpSPYALER_ | S;S | 1480;1493 | | | 1.05682 | |
| Rabep2 | Q91WG2 | _QPASLHGpSTELLPLSR_ | S | 180 | | | 1.01825 | |

Sup. Table 1 - The 117 DPPs, phosphopeptides and phosphosites

| Gene names | Protein code (Uniprot) | Phosphopeptides | (STY) sites | Phospho-sites | log2 fold change | | | |
|---|---|---|---|---|---|---|---|---|
| | | | | | shUHMK1 #2 | shUHMK1 #3 | UHMK1$^{WT}$ | UHMK1$^{K54R}$ |
| Rasa3 | Q60790 | _QQSEISTHpSI_ | S | 833 | | | 1.35870 | |
| Setd2 | E9Q5F9 | _QHNTSKpSPFR_ | S | 743 | | | 1.14517 | |
| Sugp1 | Q8CH02 | _DIDASPpSPLSVQDLK_ | S | 409 | | | 1.99911 | |
| ‡Suv39h2 | Q8K085 | _GSGEASSDSIDHpSPAK_ | S | 235 | | | 1.10931 | |
| ‡Tns1 | A0A087WQS0 | _HLGGSGSVVPGpSPSLDR_ | S | 1447 | | | 1.23857 | |
| Ubl7 | Q91W67 | _DMPGGFLFDGLpSDDEDDFHPSTR_ | S | 230 | | | 1.30980 | |
| Wnk1 | P83741 | _GTEDGSGSPHpSPPHLCSK_ | S | 2027 | | | 1.09522 | |
| Ythdf2 | Q91YT7 | _DGLNDDDFEPYLpSPQAR_ | S | 39 | | | 1.23362 | |
| Zfml | A0A0N4SV80 | _QSSVTQVTEQpSPK_ | S | 128 | | | 1.21899 | |
| Zswim8 | Q3UHH1 | _HTGMASIDSSAPETTSDSpSPTLSR_ | S | 1158 | | | 1.02638 | |
| **Proteins differentially phosphorylated solely in UHMK1$^{K54R}$** | | | | | | | | |
| Carhsp1 | Q9CR86 | _TRTFpSATVR_ | S | 53 | | | | -1.06353 |
| Cux1 | H3BK24;H3BLS0 | _RPpSSLQSLFGLPEAAGAR_ | S | 1456 | | | | -1.21232 |
| Eef1d | Q80T06;A0A0R4J1L2 | _ATAPQTQHVpSPMRQVEPPTKK_ | S | 133 | | | | -1.44256 |
| Eif2b5 | Q8CHW4 | _AGpSPQLDDIRVFQNEVLGTLQR_ | S | 540 | | | | -1.93408 |
| Eif2s2 | Q99L45 | _(ac)pSGDEMIFDPTMSKK_ | S | 2 | | | | -1.42880 |
| Pbdc1 | Q9D0B6 | _GADpSGGEKEEGANREGEK_ | S | 184 | | | | -1.64152 |
| Prpf38b | Q80SY5 | _pSIDRGLDR_ | S | 321 | | | | -1.31779 |
| Thrap3 | Q569Z6 | _ERpSTEKTEK_ | S | 750 | | | | -1.39575 |
| Zc3h18 - I | H3BIW0 | _ApSDLEEEENATR_ | S | 45 | | | | -1.09051 |
| Zc3h18 - II | H3BIW0 | _KANLSPDRGpSR_ | S | 854 | | | | 1.02756 |

‡ Phosphosites not registered in the databases PhosphositePlus and PHOSIDA (mouse).
Different phosphopeptides from the same protein are labeled with roman numbers.

Sup. Table 2 – Biological Process (BP) terms from Gene Ontology (GO) enrichment analysis of the putative UHMK1 substrates performed in PANTHER

| **Term** | **Number of proteins** | **Fold enrichment** | **raw P-value** | **FDR** |
|---|---|---|---|---|
| Maturation of LSU-rRNA from tricistronic rrna transcript (SSU-rRNA. 5.8S rRNA. LSU- rRNA) (GO:0000463) | 3 | 35.85 | 1.25E-04 | 3.46E-02 |
| RNA processing (GO:0006396) | 19 | 4.74 | 2.32E-08 | 4.58E-05 |
| Gene expression (GO:0010467) | 30 | 3.26 | 6.33E-09 | 2.50E-05 |
| Macromolecule metabolic process (GO:0043170) | 51 | 1.88 | 1.34E-06 | 1.24E-03 |
| Organic substance metabolic process (GO:0071704) | 58 | 1.62 | 1.81E-05 | 9.85E-03 |
| Metabolic process (GO:0008152) | 59 | 1.53 | 9.70E-05 | 3.06E-02 |
| RNA metabolic process (GO:0016070) | 24 | 3.81 | 1.58E-08 | 4.14E-05 |
| Nucleic acid metabolic process (GO:0090304) | 29 | 3.19 | 1.89E-08 | 4.25E-05 |
| Nucleobase-containing compound metabolic process (GO:0006139) | 30 | 2.65 | 5.55E-07 | 6.26E-04 |
| Heterocycle metabolic process (GO:0046483) | 31 | 2.58 | 6.40E-07 | 6.73E-04 |
| Cellular metabolic process (GO:0044237) | 57 | 1.70 | 4.80E-06 | 3.79E-03 |
| Cellular process (GO:0009987) | 103 | 1.31 | 1.56E-07 | 2.74E-04 |
| Primary metabolic process (GO:0044238) | 56 | 1.69 | 7.19E-06 | 5.16E-03 |
| Cellular aromatic compound metabolic process (GO:0006725) | 31 | 2.49 | 1.27E-06 | 1.25E-03 |
| Cellular nitrogen compound metabolic process (GO:0034641) | 39 | 2.69 | 3.44E-09 | 1.81E-05 |
| Nitrogen compound metabolic process (GO:0006807) | 56 | 1.85 | 3.42E-07 | 4.15E-04 |
| Organic cyclic compound metabolic process (GO:1901360) | 32 | 2.36 | 3.08E-06 | 2.56E-03 |
| Cellular component biogenesis (GO:0044085) | 26 | 2.21 | 1.39E-04 | 3.73E-02 |
| Cellular component organization or biogenesis (GO:0071840) | 48 | 1.73 | 4.12E-05 | 1.58E-02 |
| Neurotransmitter receptor transport to postsynaptic membrane (GO:0098969) | 3 | 31.87 | 1.70E-04 | 4.14E-02 |
| Neurotransmitter receptor transport to plasma membrane (GO:0098877) | 3 | 30.19 | 1.97E-04 | 4.63E-02 |
| Establishment of protein localization to postsynaptic membrane (GO:1903540) | 3 | 30.19 | 1.97E-04 | 4.56E-02 |
| Translational initiation (GO:0006413) | 5 | 17.38 | 1.52E-05 | 9.22E-03 |
| Translation (GO:0006412) | 8 | 5.03 | 2.31E-04 | 4.85E-02 |
| Amide biosynthetic process (GO:0043604) | 10 | 4.51 | 9.14E-05 | 2.94E-02 |
| Cellular biosynthetic process (GO:0044249) | 23 | 2.23 | 2.33E-04 | 4.84E-02 |
| Organonitrogen compound biosynthetic process (GO:1901566) | 16 | 2.84 | 1.69E-04 | 4.17E-02 |
| Cellular macromolecule biosynthetic process (GO:0034645) | 17 | 2.82 | 1.16E-04 | 3.39E-02 |
| Macromolecule biosynthetic process (GO:0009059) | 17 | 2.78 | 1.35E-04 | 3.68E-02 |
| Positive regulation of cell cycle G1/S phase transition (GO:1902808) | 4 | 15.93 | 1.55E-04 | 4.02E-02 |
| Positive regulation of cell cycle phase transition (GO:1901989) | 5 | 9.76 | 2.04E-04 | 4.59E-02 |
| Regulation of cell cycle (GO:0051726) | 9 | 4.87 | 1.17E-04 | 3.37E-02 |
| Regulation of cellular process (GO:0050794) | 80 | 1.34 | 1.63E-04 | 4.08E-02 |

Sup. Table 2 – Biological Process (BP) terms from Gene Ontology (GO) enrichment analysis of the putative UHMK1 substrates performed in PANTHER (To be continued)

| Term | Number of proteins | Fold enrichment | raw P-value | FDR |
|---|---|---|---|---|
| Regulation of biological process (GO:0050789) | 82 | 1.31 | 2.30E-04 | 4.91E-02 |
| Positive regulation of cell cycle (GO:0045787) | 9 | 4.87 | 1.17E-04 | 3.37E-02 |
| Positive regulation of cellular process (GO:0048522) | 51 | 1.72 | 2.33E-05 | 1.05E-02 |
| Positive regulation of biological process (GO:0048518) | 54 | 1.68 | 1.59E-05 | 9.31E-03 |
| Regulation of microtubule polymerization (GO:0031113) | 4 | 14.43 | 2.22E-04 | 4.80E-02 |
| Regulation of microtubule polymerization or depolymerization (GO:0031110) | 5 | 10.51 | 1.46E-04 | 3.85E-02 |
| Regulation of microtubule cytoskeleton organization (GO:0070507) | 6 | 7.50 | 1.86E-04 | 4.45E-02 |
| Regulation of cytoskeleton organization (GO:0051493) | 11 | 3.98 | 1.18E-04 | 3.34E-02 |
| Regulation of protein polymerization (GO:0032271) | 8 | 6.71 | 3.32E-05 | 1.31E-02 |
| Regulation of protein-containing complex assembly (GO:0043254) | 11 | 4.79 | 2.35E-05 | 1.03E-02 |
| Regulation of cellular component biogenesis (GO:0044087) | 17 | 3.39 | 1.20E-05 | 7.60E-03 |
| Positive regulation of protein localization to cell periphery (GO:1904377) | 5 | 13.86 | 4.23E-05 | 1.59E-02 |
| Positive regulation of cellular protein localization (GO:1903829) | 8 | 5.73 | 9.72E-05 | 3.01E-02 |
| Regulation of protein localization to cell periphery (GO:1904375) | 6 | 8.44 | 1.00E-04 | 3.04E-02 |
| Negative regulation of protein polymerization (GO:0032272) | 5 | 12.75 | 6.15E-05 | 2.16E-02 |
| Negative regulation of protein-containing complex assembly (GO:0031333) | 6 | 8.25 | 1.12E-04 | 3.35E-02 |
| Regulation of mRNA splicing. via spliceosome (GO:0048024) | 6 | 9.89 | 4.31E-05 | 1.58E-02 |
| Regulation of mRNA processing (GO:0050684) | 7 | 8.69 | 2.16E-05 | 1.00E-02 |
| Regulation of mRNA metabolic process (GO:1903311) | 11 | 7.94 | 2.13E-07 | 3.36E-04 |
| Regulation of RNA splicing (GO:0043484) | 7 | 8.47 | 2.53E-05 | 1.05E-02 |
| Positive regulation of translation (GO:0045727) | 6 | 8.96 | 7.27E-05 | 2.49E-02 |
| Positive regulation of cellular amide metabolic process (GO:0034250) | 6 | 7.26 | 2.20E-04 | 4.89E-02 |
| Posttranscriptional regulation of gene expression (GO:0010608) | 11 | 4.76 | 2.49E-05 | 1.06E-02 |
| Regulation of mRNA catabolic process (GO:0061013) | 6 | 8.96 | 7.27E-05 | 2.44E-02 |
| mRNA splicing. via spliceosome (GO:0000398) | 8 | 7.25 | 1.95E-05 | 9.92E-03 |
| mRNA processing (GO:0006397) | 15 | 6.88 | 7.35E-09 | 2.32E-05 |
| mRNA metabolic process (GO:0016071) | 18 | 6.43 | 5.63E-10 | 4.44E-06 |
| RNA splicing. via transesterification reactions with bulged adenosine as nucleophile (GO:0000377) | 8 | 7.25 | 1.95E-05 | 9.61E-03 |
| RNA splicing. via transesterification reactions (GO:0000375) | 8 | 7.25 | 1.95E-05 | 9.32E-03 |
| RNA splicing (GO:0008380) | 15 | 8.69 | 3.44E-10 | 5.42E-06 |
| Mitotic cell cycle process (GO:1903047) | 12 | 4.93 | 7.34E-06 | 5.04E-03 |
| Mitotic cell cycle (GO:0000278) | 13 | 4.64 | 5.75E-06 | 4.32E-03 |

Sup. Table 2 – Biological Process (BP) terms from Gene Ontology (GO) enrichment analysis of the putative UHMK1 substrates performed in PANTHER (Conclusion)

| Term | Number of proteins | Fold enrichment | raw P-value | FDR |
|---|---|---|---|---|
| Cell cycle (GO:0007049) | 22 | 3.49 | 3.12E-07 | 4.10E-04 |
| Cell cycle process (GO:0022402) | 18 | 4.30 | 2.30E-07 | 3.30E-04 |
| Cell division (GO:0051301) | 12 | 4.72 | 1.13E-05 | 7.44E-03 |
| Microtubule cytoskeleton organization (GO:0000226) | 12 | 4.52 | 1.74E-05 | 9.79E-03 |
| Cytoskeleton organization (GO:0007010) | 16 | 2.78 | 2.21E-04 | 4.84E-02 |
| Organelle organization (GO:0006996) | 32 | 1.94 | 1.56E-04 | 3.98E-02 |
| Cellular component organization (GO:0016043) | 46 | 1.72 | 8.63E-05 | 2.84E-02 |
| Microtubule-based process (GO:0007017) | 13 | 3.26 | 1.99E-04 | 4.55E-02 |
| Cellular protein-containing complex assembly (GO:0034622) | 13 | 3.70 | 5.74E-05 | 2.06E-02 |
| Protein-containing complex subunit organization (GO:0043933) | 18 | 3.00 | 3.16E-05 | 1.28E-02 |
| Cellular response to stress (GO:0033554) | 21 | 2.79 | 1.92E-05 | 1.01E-02 |

Highlighted in grey: the most specific BP terms within the hierarchy presented in PANTHER, depicted in Figure 4a.

Sup. Table 3 – Differentially Expressed Genes (DEGs) upon UHMK1 knockdown (UHMK1-KD *vs*. shCTRL)

| **Count** | **Gene names** | **log2 fold change** | **Adj. p-value** |
|---|---|---|---|
| 1 | *Oaz1-ps* | 0.6691 | 0.00000000 |
| 2 | *Oaz1* | 0.7167 | 0.00000000 |
| 3 | *Uhmk1* | -1.6634 | 0.00000000 |
| 4 | *Pimreg* | 0.4393 | 0.00002744 |
| 5 | *Ccl9* | -0.4246 | 0.00002744 |
| 6 | *Fam45a* | 0.6287 | 0.00007735 |
| 7 | *Cdca4* | 0.5593 | 0.00008340 |
| 8 | *Lox* | -0.4747 | 0.00019220 |
| 9 | *Gja1* | 0.4010 | 0.00068750 |
| 10 | *Map3k3* | 0.5132 | 0.00092238 |
| 11 | *Sptbn1* | -0.3553 | 0.00096137 |
| 12 | *Npr3* | -0.5548 | 0.00107712 |
| 13 | *Adh7* | 0.4975 | 0.00107712 |
| 14 | *Fads1* | 0.3154 | 0.00107712 |
| 15 | *Pcdh19* | -0.4722 | 0.00758685 |
| 16 | *Rab28* | 0.4464 | 0.01917932 |
| 17 | *Erlin2* | 0.4334 | 0.01917932 |
| 18 | *Ube2g1* | 0.3665 | 0.02933273 |
| 19 | *Wfdc21* | -1.2072 | 0.03024229 |
| 20 | *Herpud2* | 0.5569 | 0.03324926 |
| 21 | *Taldo1* | 0.1761 | 0.03394717 |
| 22 | *Ndufa8* | 0.2585 | 0.03584492 |
| 23 | *Gask1b* | -0.4354 | 0.03677648 |
| 24 | *Prrx1* | -0.3072 | 0.03803463 |
| 25 | *Memo1* | -0.2756 | 0.03803463 |
| 26 | *Mdh1* | 0.2472 | 0.03803463 |
| 27 | *Tpt1* | -0.1810 | 0.03944166 |
| 28 | *Pkia* | -0.6053 | 0.04036702 |
| 29 | *Kank3* | -0.3181 | 0.04300979 |
| 30 | *Dmpk* | 0.7275 | 0.04677956 |
| 31 | *Far1* | -0.2162 | 0.04677956 |
| 32 | *Tpt1-ps6* | -0.2162 | 0.04979391 |
